# Genome-wide analysis of transcription-coupled repair reveals novel transcription events in *Caenorhabditis elegans*

**DOI:** 10.1101/2023.10.12.562083

**Authors:** Cansu Kose, Laura A. Lindsey-Boltz, Aziz Sancar, Yuchao Jiang

## Abstract

Bulky DNA adducts such as those induced by ultraviolet light are removed from the genomes of multicellular organisms by nucleotide excision repair, which occurs through two distinct mechanisms, global repair, requiring the DNA damage recognition-factor XPC (xeroderma pigmentosum complementation group C), and transcription-coupled repair (TCR), which does not. TCR is initiated when elongating RNA polymerase II encounters DNA damage, and thus analysis of genome-wide excision repair in XPC-mutants only repairing by TCR provides a unique opportunity to map transcription events missed by methods dependent on capturing RNA transcription products and thus limited by their stability and/or modifications (5’-capping or 3’- polyadenylation). Here, we have performed the eXcision Repair-sequencing (XR-seq) in the model organism *Caenorhabditis elegans* to generate genome-wide repair maps from a wild-type strain with normal excision repair, a strain lacking TCR (*csb-1*), or one that only repairs by TCR (*xpc- 1*). Analysis of the intersections between the *xpc-1* XR-seq repair maps with RNA-mapping datasets (RNA-seq, long- and short-capped RNA-seq) reveal previously unrecognized sites of transcription and further enhance our understanding of the genome of this important model organism.

## INTRODUCTION

Transcription of eukaryotic genomes by RNA polymerase II (RNAPII) produces both protein-coding mRNAs and diverse non-coding RNAs (ncRNAs), including enhancer RNAs (eRNAs), long intergenic non-coding RNAs (lincRNAs), and Piwi-interacting RNAs (piRNAs) [1]. Most ncRNAs are rapidly degraded making them difficult to detect, and therefore, likely to have not been fully mapped [2]. However, proper mapping of transient RNAs is an important first step towards understanding the function of these ncRNAs [3]. Many methods have been developed to capture and sequence RNAPII transcripts including those that harness RNA capture through their modifications, such as 3’-polyadenylation (poly(A)), used in RNA-seq [4], and 5’-capping used in capped RNA-seq [5, 6]. Incorporation of an RNA size-selection step in the later technique to specifically capture short- or long-capped RNAs of less than 100 nucleotides (nt) or greater than 200 nt, respectively, has been beneficial in identifying different classes of ncRNAs and revealed many novel sites of transcription [5, 6]. Here, we describe a unique way of identifying RNAPII transcription, which is independent of capturing the RNA products, but instead, harnesses the mechanistic properties of nucleotide excision repair and a sensitive method for sequencing whole-genome excision repair events called XR-seq for e<u>X</u>cision <u>R</u>epair<u>-seq</u>uencing [7] **(Fig 1A)**.

**Fig 1.**
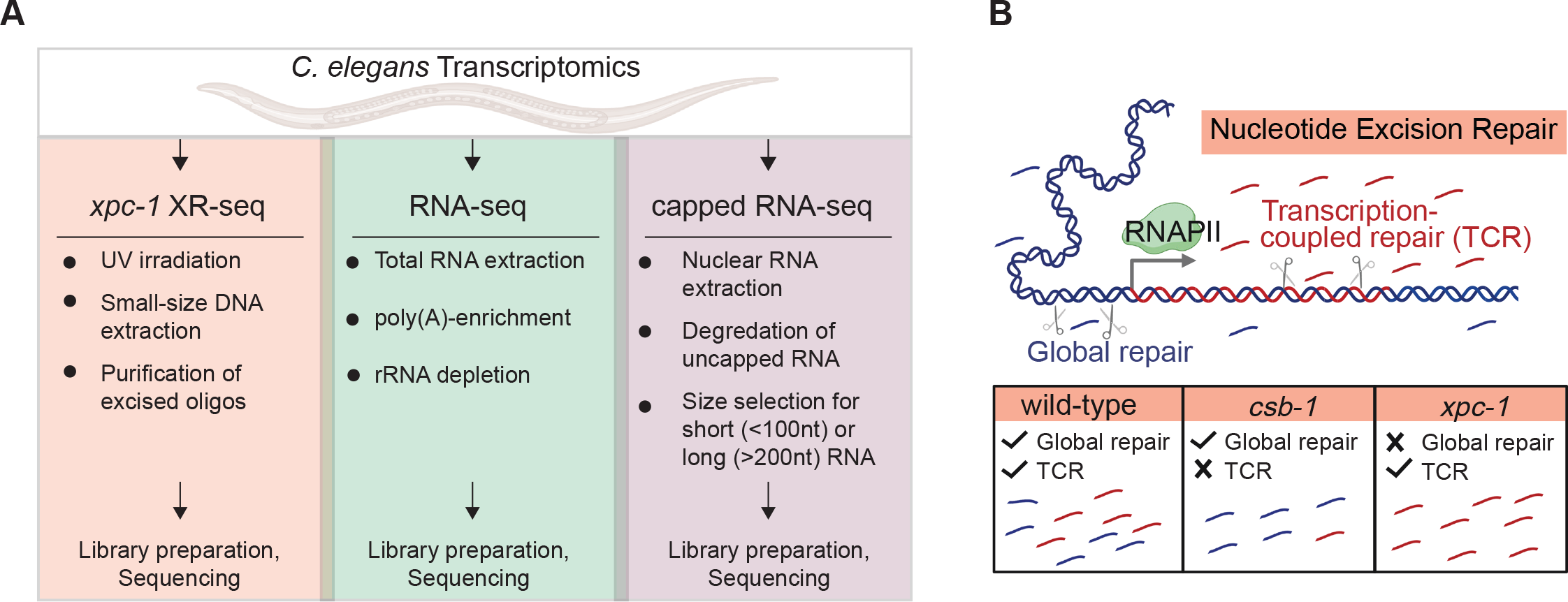
Overview of Study Design. (A) The illustration highlights key properties of the three comparative transcriptomic techniques (XR-seq, RNA-seq, capped RNA-seq) analyzed in this study for their capacity to identify genome-wide transcription in *C. elegans*. **(B)** Nucleotide excision repair removes DNA damage through two different mechanisms: global repair and transcription-coupled repair (TCR). Global repair depends on the XPC protein and occurs throughout the genome, whereas TCR is independent of XPC and only occurs when elongating RNA polymerase II encounters damage during transcription and recruits the CSB protein. This study uses XR-seq to map nucleotide excision repair at single-nucleotide resolution throughout the whole-genome in three strains of *C. elegans*: wild-type, *csb-1*, and *xpc-1*. Because *xpc-1* worms lack global repair, analysis of XR-seq data from this strain provides a unique opportunity to map transcription genome-wide independent of RNA capture.

In eukaryotes, nucleotide excision repair removes a wide range of helix-distorting DNA lesions from the genome, including UV-induced cyclobutane pyrimidine dimer (CPDs), by concerted dual incision of the phosphodiester bonds bracketing the lesion at a somewhat precise distance from the damage (∼19 nt 5′ and ∼6 nt 3′ to the dimer) to generate ∼27-nt damage-containing oligonucleotide excision products [8, 9]. Nucleotide excision repair is carried out in most eukaryotes by the six excision repair factors XPA through XPG, originally identified by complementation assays using cells from Xeroderma Pigmentosum (XP) patients, which exhibit a hereditary condition characterized by extreme sun-sensitivity and skin cancer incidence [10]. *Caenorhabditis elegans (C. elegans)* have homologs of the human XP excision repair factors except for XPE (DDB2) [11]. In addition to these factors, two additional proteins, which are also conserved in *C. elegans* [12], CSA and CSB, were subsequently identified in patients with a related human genetic disorder called Cockayne Syndrome (CS), exhibiting photosensitivity resulting from deficient transcription-coupled repair (TCR), which is defined as excision repair of DNA damage specifically within the transcribed strand of actively transcribed DNA [13]. Nucleotide excision repair occurs by two mechanistically distinct pathways: global repair, that depends on XPA through XPG, and TCR, that depends on these same factors excluding XPC [10]. TCR is initiated when damage in the template strand is encountered by elongating RNAPII, which subsequently recruits CSB, CSA, and additional factors. *C. elegans* have been shown to repair UV-induced DNA damage by both global repair and TCR pathways [11, 12, 14–17].

We conducted the current study in the *C. elegans* worm model organism because of its nearly completely annotated nuclear genome, which is approximately 1/30 the size of the human genome, and because of the availability of a range of omics data. To avoid complications of conducting experiments on mixed populations of whole animals at different developmental stages, many *C. elegans* study designs employ collecting L1-stage larvae state of developmental arrest and uniformly stimulating them into progression upon feeding in order to gather sizable cohorts of animals at a singular developmental phase. *C. elegans* studies of DNA repair have also been performed using L1-stage worms [11, 12, 14–17], and there are a multitude of available omics data sets examining epigenetic markers, chromatin states, and RNA expression at this stage [5, 6, 18], so we chose to conduct the current study at this stage as well. We performed XR-seq and RNA-seq in three previously characterized strains of *C. elegans*: wild-type (WT) exhibiting both global repair and TCR, *csb-1*, which only repairs by global repair, and *xpc-1*, which only has TCR **(Fig 1B)**. We provide evidence demonstrating the utility of *xpc-1* XR-seq data set for detecting RNAPII transcription and identifying new transcripts. The integration of epigenetic markers, chromatin states, and ncRNA annotations including eRNAs, lincRNAs, and piRNAs all support the robust detection of intergenic RNAPII transcription by *xpc-1* XR-seq. Overall, our results provide a comprehensive view of the transcription-coupled repair landscape in *C. elegans*, highlighting its potential contribution to our understanding of DNA repair mechanisms and non-coding RNA biology.

## RESULTS

### XR-seq repair maps of UV-induced DNA damage in wild-type, csb-1, and xpc-1 strains of C. elegans

The XR-seq next generation sequencing method (Fig S1) was developed to capture and identify DNA damage-containing excised oligomers to map repair at single-nucleotide resolution throughout the human genome [19]. Recently we modified the method to analyze excision repair of UV-induced CPD photoproducts in *C. elegans* and demonstrated that excision repair in *xpa-1* mutants was near background (that of unirradiated WT worms) [20]. Here we have extended our study to include two additional previously characterized repair-deficient *C. elegans* strains, *xpc-1* and *csb-1* [12, 17, 21] (see **S1 Table** for detailed sample information). **Fig 2A** shows a representative Integrative Genomics Viewer (IGV) screenshot of a 5.2 kilobase (kb) region of the genome containing two genes in opposite orientations illustrating levels of transcription as measured by RNA-seq (top two rows) and repair as measured by XR-seq (remaining rows) in WT, *csb-1* and *xpc-1* strains. Reads are mapped to the two strands of the genome as shown, and for the *vbh-1* gene, the transcribed strand (TS) is the + strand and for the *mrpl-17* gene, the TS is the – strand. It is noteworthy that the reads acquired from XR-seq align to the template strand and are complementary to those obtained from RNA-seq, which align to the coding strand of the gene. The results illustrate preferential repair of the TS due to TCR in both WT and *xpc-1*, and there is no strand preference observed in the *csb-1* strain. Quantitatively, we used the ratio of read counts from the TS to those from both the TS and non-transcribed strand (NTS) as a proxy for TCR, with genome-wide results shown in **S2 Table**. The percentage of TS/(TS + NTS) for the *vbh-1* and *mrpl- 17* genes are, respectively, 78% and 77% in WT, which has both global and TCR; 99% and 94% in *xpc-1*, which only has TCR; and 59% and 43% in *csb-1*, which only has global repair. As previously shown, the unirradiated control (WT no UV) results in extremely low read-numbers (0.003% of UV-irradiated WT) that are not specific [20].

**Fig 2.**
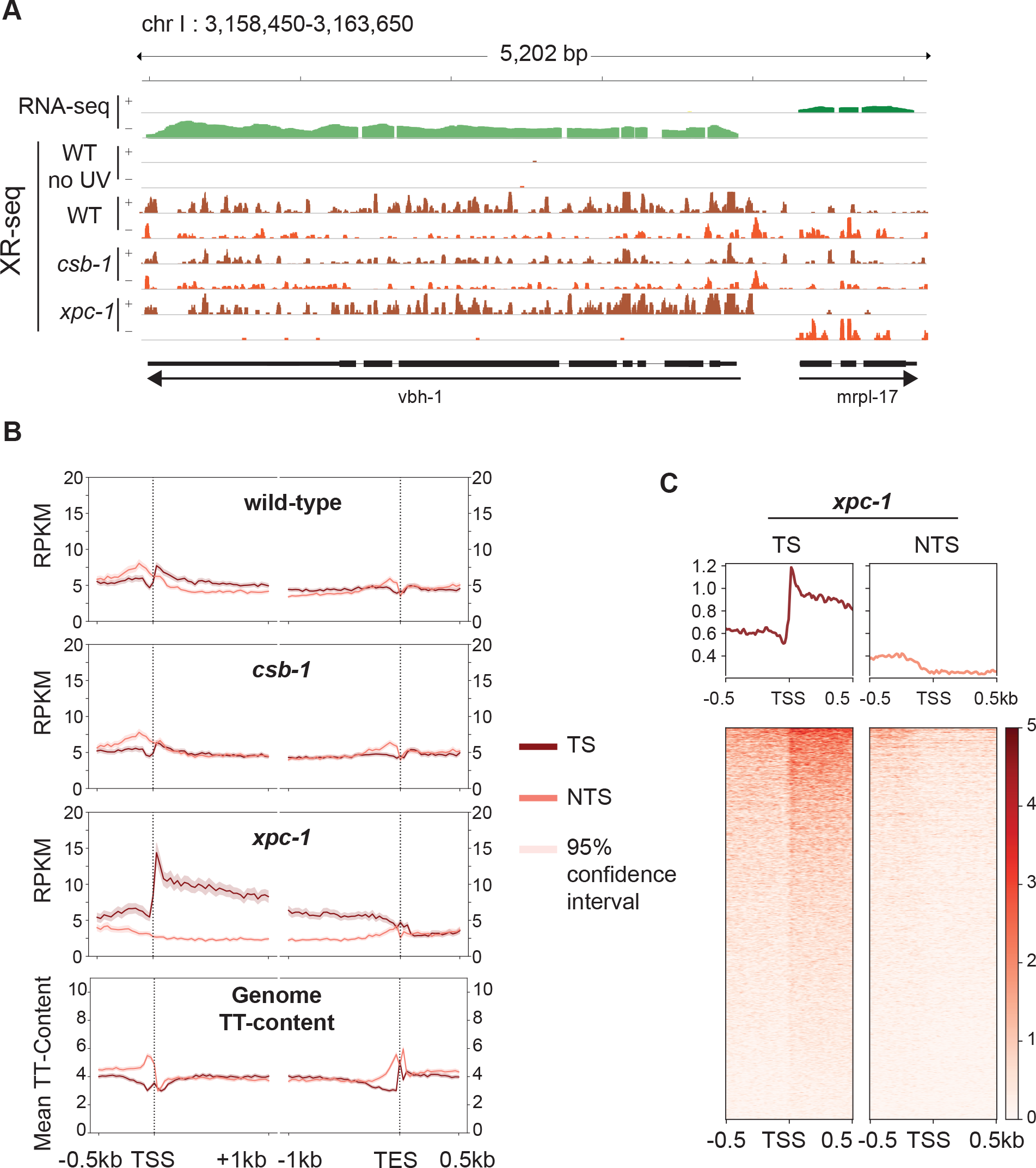
Detection of Transcription-Coupled Repair by XR-seq. (A) Browser view of the distribution of *C. elegans* high throughput sequencing reads separated by strand over a representative 5.2 kb region from chromosome I. RNA-seq reads (green) from wild-type (WT) worms is shown on top to illustrate the opposite direction of transcription of the genes *vbh-1* and *mrpl-17*. The strand distribution of XR-seq reads (orange) 1 hour after UV clearly demonstrates the occurrence of transcription-coupled repair within the body of both genes in WT and *xpc-1* worms but absent in *csb-1*. **(B)** To analyze transcription-coupled repair genome-wide, XR-seq reads on transcribed strand (TS) and non-transcribed strand (NTS) in the indicated strains at 1 hour repair time is plotted with mean reads per kilobase per million mapped reads (RPKM) (y-axis) along the 500 bp upstream and 1 kb downstream of transcription start site (TSS), and 1 kb upstream and 500 bp downstream of transcription end site (TES) (x-axis) for 2,142 genes selected for length > 2 kb and no overlaps with a distance of at least 500 bp between genes. The TT-distribution, as mean TT content (y-axis) was determined for the same gene set and is plotted at the bottom as a measure of expected DNA damage sites. **(C)** Profile plots and heatmaps of TS and NTS XR-seq reads from the *xpc-1* strain at 1 hour repair time spanning the best represented half of 16,588 TSSs > 1 kb apart indicate divergent transcription at promoters.

The XR-seq data were then analyzed to assess repair on the TS and NTS of all non- overlapping genes greater than 2 kb in length (**Fig 2B**). Again, such analysis clearly illustrates the presence of TCR in the WT strain (top), which is partially masked due to global repair products mapping to both strands. There was no observed strand difference in repair within gene bodies in the *csb-1* mutant (middle), which lacks TCR. Notably, the differences observed upstream of the transcription start site (TSS) and transcription end site (TES) can be attributed to TT-content of these regions of the genome (bottom) resulting in different levels of DNA damage in these areas since CPDs are primarily formed at TTs and the majority of XR-seq reads contain this dinucleotide ∼6 nt from the 3’-end (**S2, S3A Fig)**. Within gene bodies the TT-content does not significantly vary with gene length or between strands in *C. elegans* (**S4 Fig)**. In stark contrast to the *csb-1* mutant, the majority of repair events map to the transcribed strand in the *xpc-1* mutant where TCR is the only functional excision repair pathway **(Fig 2B)**. As previously seen in humans and other organisms [19, 22–25], XR-seq reads peak at the 5’-end of genes and decrease toward the 3’-end, which is consistent with the TCR model proposed by Chiou *et al.* [26], and the skewed pattern gradually diminishes as the repair process proceeds over time (**S3B Fig)**. The 5’-peak of repair on the TS is not unique to L1-stage worms as this pattern is also observed in a mixed population of worms (**S3C Fig)**.

XR-seq analysis in human XP-C cells revealed pronounced CPD repair on the non- template strand upstream of the TSS due to divergent transcription at promoters [19, 25]. The *C. elegans* XR-seq analysis shown in **Fig 2B** does not exhibit this pattern even though anti-sense transcription at promoters has been reported in worms [6]. Therefore, we further analyzed repair in the region of a greater number of TSSs (all TSSs that are at least 1kb apart) at an individual basis as visualized in the heatmaps shown in **Fig 2C** and **S5 Fig**. With this analysis, we were able to detect anti-sense transcription (enrichment on NTS upstream of the TSS) in a subset of genes. The TS upstream of the TSS has much higher read-count, likely due to extensive transcription of upstream eRNAs, which has been reported to be prevalent in *C. elegans* and occur in the direction of the downstream gene in 90% of cases [6].

We next sought to identify genes that exhibit significantly differential and dynamic repair using time-series *xpc-1* XR-seq data collected at 5min, 1h, 8h, 16h, 24h, and 48h after UV- irradiation (see Materials and Methods for details). We identified 121 genes exhibiting significant dynamic repair across timepoints **(S6A Fig)** and performed gene ontology (GO) analysis of biological processes **(S6B Fig)** and cellular components **(S6C Fig)**. While investigation of gene- specific excision repair has been extensively explored across various model organisms [23, 24, 27–33] and across different timepoints [16, 34, 35], our current investigation centers on the domain of intergenic transcription-coupled repair and its juxtaposition with transcriptional events detectable by RNA-seq and capped RNA-seq.

### Transcription-coupled repair measured by XR-seq in xpc-1 C. elegans serves as an RNA-independent proxy for transcription

Since the *xpc-1* worm mutant lacks global repair, the XR-seq reads from this strain can serve as a unique measure of RNAPII transcription independent of capturing the RNA product. **Fig 3A** shows an IGV screenshot of a 27 kb region of the genome illustrating levels of transcription as measured by RNA-seq in WT and *xpc-1* strains, long- and short-capped RNA-seq from the WT strain, and XR-seq from the *xpc-1* strain. This representative region shows genes on either side of an intergenic region (defined as a region at least 2 kb away from an annotated gene). RNA-seq reads (top) can be seen in the areas of the annotated RefSeq Genes consistent with polyadenylated protein-coding mRNA transcripts. We do not observe obvious differences in the RNA-seq data from the two different worm strains (WT and *xpc-1*) (Spearman correlation coefficient, r=0.94). The long-capped RNA-seq reads, which do not require poly(A) for capture, are seen in these same areas of protein-coding transcripts and are also seen in the intergenic region. This is consistent with previous reports demonstrating that this technique is useful for detecting non-coding RNAs [6, 18]. Similarly, short-capped RNA-seq reads have been reported to effectively map areas of transcription initiation, of which there are many in this screenshot. There are *xpc-1* XR-seq reads (bottom) throughout this highly transcribed 27 kb area of the genome, including the intergenic region, which illustrates the potential value of using the data set as an RNA-independent proxy for transcription.

**Fig 3.**
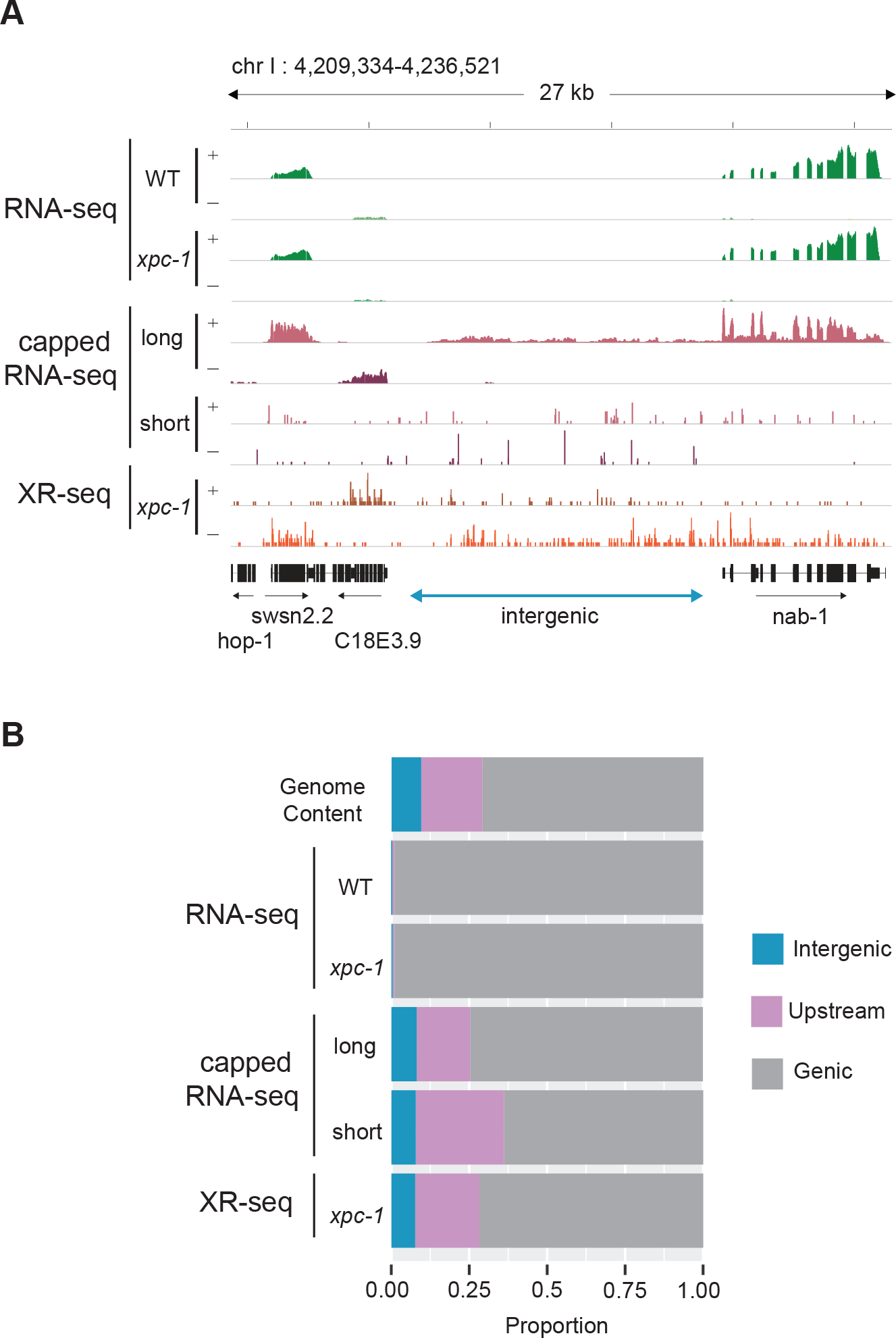
Transcription-Coupled Repair Reveals Transcription in Intergenic Regions. (A) Browser view of stranded read distribution from WT and *xpc-1* RNA-seq (green), capped RNA- seq (pink), and *xpc-1* XR-seq (orange) at 1 hour repair time over a representative 27 kb region from chromosome I. Both capped RNA-seq and XR-seq methods provide comprehensive coverage of the entire window, encompassing both genic and intergenic regions, in contrast to RNA-seq which only captures polyadenylated mRNAs. **(B)** Bar graphs depict the genome-wide distribution of reads obtained from the different sequencing methods, including WT and *xpc-1* RNA-seq, long- capped RNA-seq, short-capped RNA-seq, and *xpc-1* XR-seq at 1 hour repair time. Notably, both XR-seq and capped RNA-seq techniques reveal transcription events occurring outside of the defined genic regions (see Materials and Methods for details).

We compared the genome-wide distribution of the reads obtained from the different sequencing methods **(Fig 3B, S7A Fig)**. For this analysis, the genome was systematically divided into three distinct categories: intergenic regions, regions within 2 kb upstream of TSSs, and genic regions. Notably, both *xpc-1* XR-seq and capped RNA-seq techniques reveal a large proportion of transcription events occurring outside of genic regions. This analysis reveals a noteworthy distinction when comparing RNA-seq, capped RNA-seq, and XR-seq. In contrast to RNA-seq, both capped RNA-seq and *xpc-1* XR-seq generate a significantly higher number of reads that map to intergenic regions and regions located within 2 kilobases upstream of TSS. This observation underscores the capability of these methods to capture transcriptional activity in these specific genomic locations. Similarly, our investigations demonstrate a high degree of concordance between genome-wide signals obtained from XR-seq and those derived from short and long- capped RNA-seq. Conversely, there is a near-zero correlation coefficient when comparing RNA-seq to the capped RNA-seq and XR-seq datasets **(S8 Fig)**.

### Epigenetic markers and chromatin states validate the intergenic transcription detected by xpc- 1 XR-seq

Expanding our investigation further, we incorporated annotation of chromatin states of *C. elegans*. As illustrated in **Fig 4A** and **S7B Fig**, our analysis of chromatin states has unveiled intriguing distinctions among the different sequencing methods. Notably, when we examine the distribution of chromatin states, RNA-seq appears to predominantly align with 5’ proximal regions, gene bodies, and exons. However, it displays relatively lower read counts in categories associated with retrotransposons, pseudogenes, and tissue-specific regions. In stark contrast, both capped RNA-seq and XR-seq exhibit notably similar chromatin state patterns, although some nuanced differences do exist between the two. A closer examination demonstrates that both short-capped RNA-seq and long-capped RNA-seq reveal genic and intergenic transcription, including intergenic enhancers. Short-capped RNA-seq indicates shorter transcripts, corresponding to transcription initiation events and enhancers shorter than 100 base pairs. In contrast, long-capped RNA-seq captures longer transcripts within the nucleus, encompassing both pre-mature and mature RNAs. These longer transcripts relate to transcription elongation, enhancer regions, and tissue-specific transcription. Furthermore, categories that align with XR-seq encompass a combination of short- and long-capped RNA-seq signals, indicating the concordance between XR-seq and capped RNA-seq in capturing transcriptional events.

**Fig 4.**
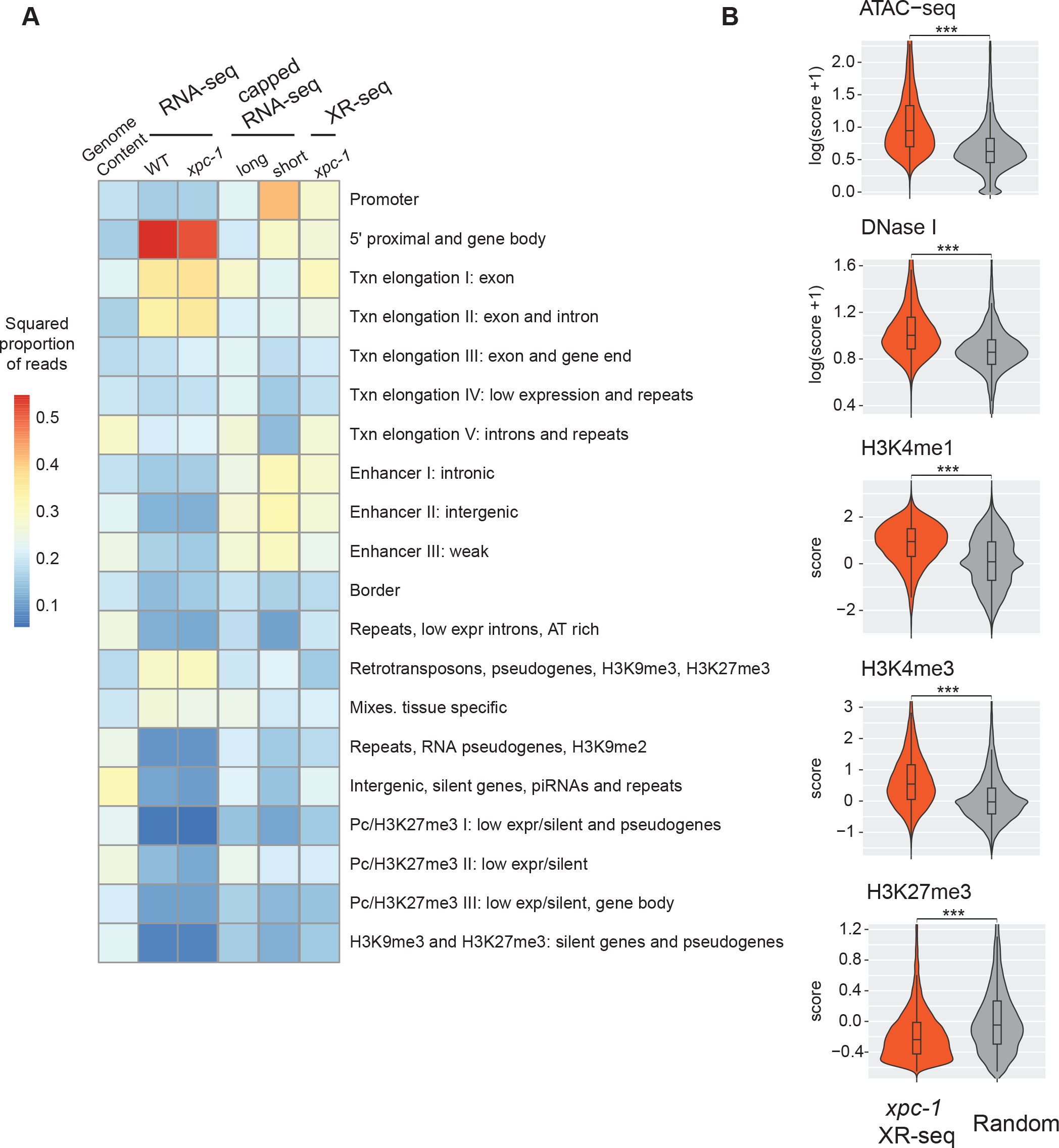
The Transcription-Coupled Repair in Intergenic Regions Detected by *xpc-1* XR-seq is Supported by Epigenomic Signatures. (A) Reads from XR-seq at 1 hour repair time, capped RNA-seq, and RNA-seq were analyzed for overlap with genomic intervals corresponding to 20 distinct predicted chromatin states in *C. elegans*. The proportion of reads was computed for each of the annotated chromatin states and the square root of the proportion is visualized as a heatmap. **(B)** Examination of intergenic XR-seq reads, which are undetectable by RNA-seq, in association with ATAC-seq, DNase-seq, H3K4me1, H3K4me3, and H3K27me3 peaks. XR-seq reads exhibit a strong correlation with active transcription markers, in contrast to the repressive marker H4K27me3, when compared to randomly selected genomic regions. All p-values obtained are highly significant (< 2.2e-16) according to nonparametric Wilcoxon rank sum tests.

In our comprehensive analysis of transcribed intergenic regions specifically identified by *xpc-1* XR-seq (but not detected by RNA-seq), we focused on histone markers and chromatin accessibility (**Fig 4B**). When compared to randomly selected genomic regions spanning the entire genome, the regions uniquely pinpointed by *xpc-1* XR-seq exhibited distinct epigenomic signatures. Specifically, these regions displayed significantly heightened chromatin accessibility, indicating a more open chromatin structure conducive to transcription. Additionally, we observed increased overlap with histone markers such as H3K4me1 and H3K4me3, typically associated with promoters and enhancers. Conversely, there were less reads overlapping with regions with histone marker H3K27me3, associated with gene repression. These corroborating epigenomic signatures serve as compelling evidence reaffirming the existence of intergenic transcription detected by *xpc-1* XR-seq. Furthermore, they underscore the utility of transcription-coupled repair as a proxy for uncovering previously elusive intergenic transcriptional events within the genome.

### Novel intergenic transcription identified with xpc-1 XR-seq

We next examined RNA-seq, XR-seq, and long- and short-capped RNA-seq read density specifically within three classes of annotated intergenic ncRNAs: enhancer RNAs (eRNAs) (**Fig 5A** and **S9A Fig**), long intergenic non-coding RNAs (lincRNAs) (**Fig 5B** and **S9B Fig**), and Piwi-interacting RNAs (piRNAs) (**Fig5C** and **S9C Fig**). Heatmaps (left) display normalized read counts for the individual annotated intergenic ncRNAs segregated by chromosomes and the bar graphs (right) summarize the log-normalized read counts for the class of ncRNA. Our findings reveal that the RNA-seq method shows limited ability to detect any of these intergenic ncRNA transcripts. This is likely attributed to the lack of poly(A) tailing of ncRNAs, which prevent them from being captured by the conventional RNA-seq technique. Both eRNAs and lincRNAs are very well-represented in the data obtained from *xpc-1* XR-seq and long- and short-capped RNA-seq. Interestingly, read density at piRNAs is high for both long-cap RNA seq and *xpc-1* XR-seq, but not short-capped RNA seq. The findings from the read density analysis of these three major classes of known *C. elegans* intergenic ncRNAs demonstrate the utility of mapping such transcripts with transcription-coupled repair.

**Fig 5.**
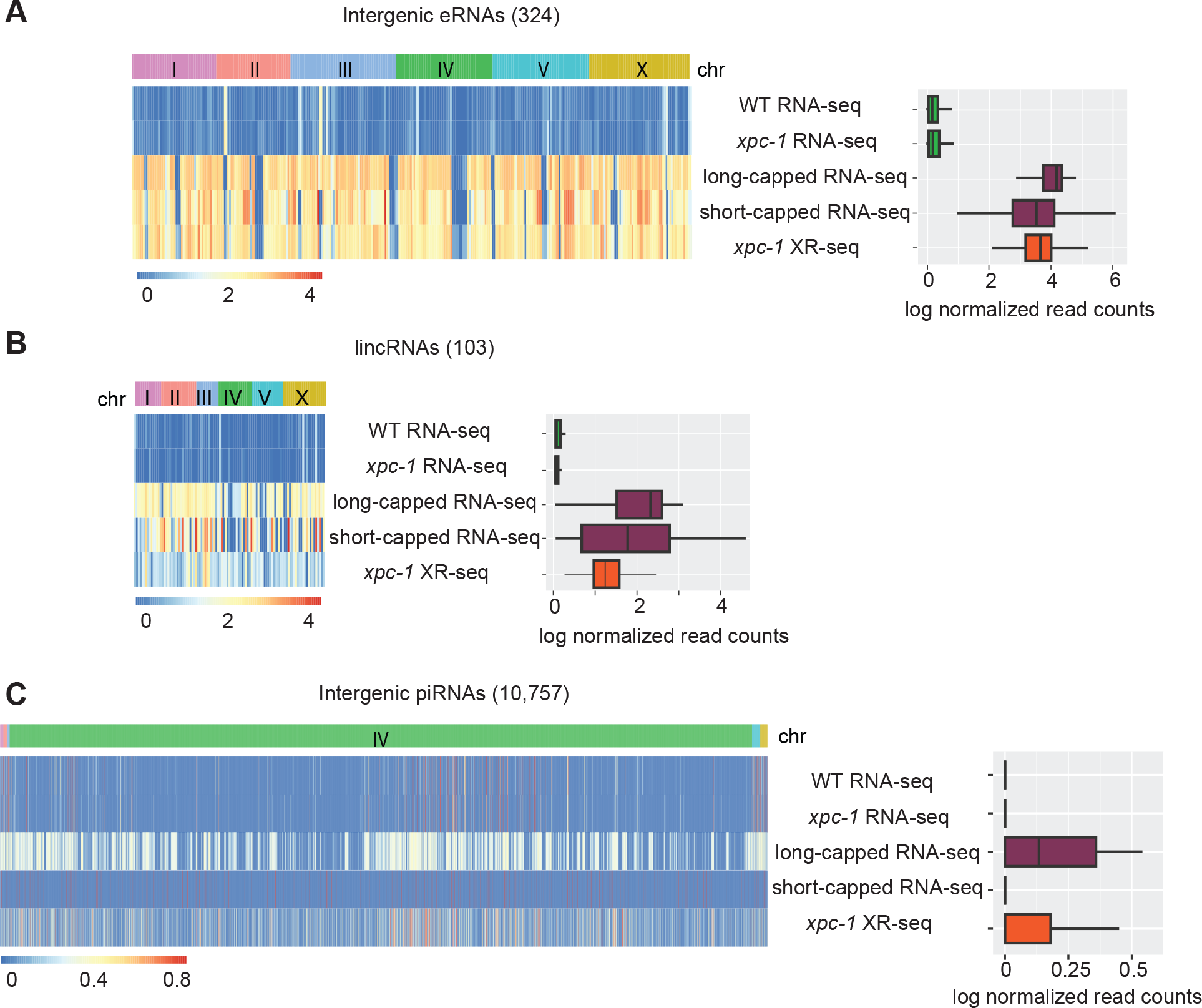
XR-seq Reveals Transcription-Coupled Repair in Intergenic eRNAs, lincRNAs, and piRNAs. (A) Heatmaps (left) display normalized RNA expression and transcription-coupled repair for intergenic enhancer RNAs (eRNAs) segregated by chromosomes. Normalization by log(x+1) was carried out, where x is library-size-adjusted read count. Bar graphs (right) represent log-normalized read counts for eRNA. Data are presented for WT and *xpc-1* RNA-seq, WT long- and short-capped RNA-seq, and time-course combined *xpc-1* XR-seq dataset (5min, 1h, 8h, 16h, 24h, and 48h). **(B, C)** Heatmaps and bar graphs as in A, for long intergenic non-coding RNAs (lincRNAs) and intergenic Piwi-interacting RNAs (piRNAs), respectively.

To assess all intergenic regions (annotated and unannotated) to determine the degree of coverage and overlap between the three methods, we divided the intergenic regions into 85,418 bins and identified those containing *xpc-1* XR-seq, RNA-seq, or capped RNA-seq reads (**S10 Fig)**. The results depicted in the Venn diagram presented in **Fig 6A** show several compelling insights. First, our analysis demonstrates that the transcription-coupled repair in the intergenic regions identified by *xpc-1* XR-seq exhibit similar coverage and remarkable concordance with capped RNA-seq, with both exhibiting ∼83% bin-coverage and 80% overlap between the two datasets. Second, as observed with the analyses above, RNA-seq has low coverage in intergenic regions relative to XR-seq and capped RNA-seq. Third, 10% of the bins contain reads unique to *xpc-1* XR-seq. Taken together, these results underscore the sensitivity of transcript-detection by *xpc-1* XR- seq.

**Fig 6.**
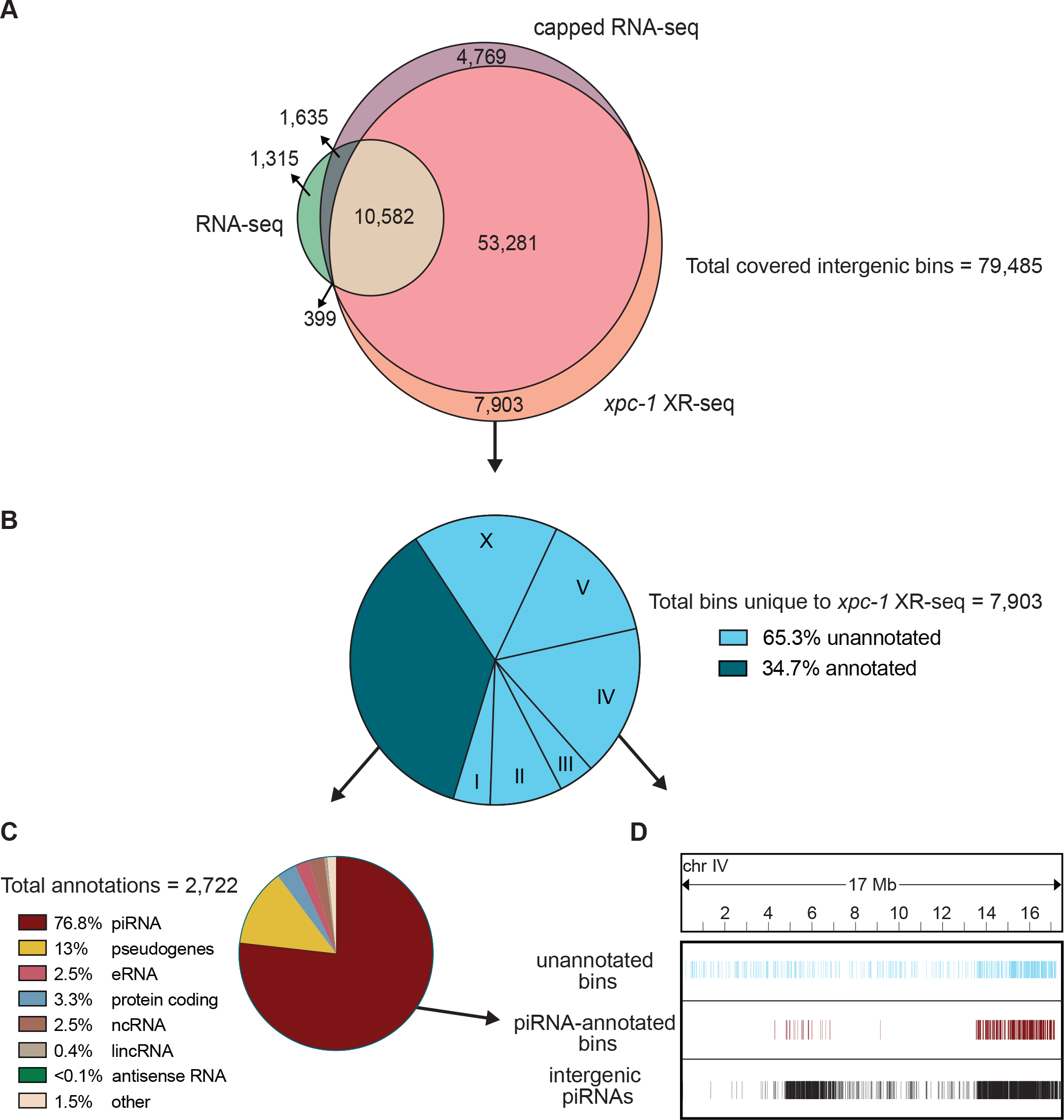
XR-seq identifies intergenic transcription-coupled repair in high concordance with intergenic transcription identified by capped RNA-seq and reveals novel sites of transcription. For 85,418 intergenic bins, we identified regions with non-zero read counts by short- or long-capped RNA-seq, RNA-seq, and time-course (5mins, 1h, 8h, 16h, 24h, and 48h repair times) *xpc-1* XR-seq. (A) Venn diagram of intergenic bins detected by capped RNA-seq, conventional RNA-seq, and *xpc-1* XR-seq. To reduce the number of call sets, we required non- zero read counts to be detected: (i) in both replicates for *xpc-1* XR-seq; (ii) in both WT and *xpc-1* RNA-seq, as they are highly correlated; and (iii) by either short-capped or long-capped RNA-seq, as they are complementary. (B) Pie chart summary of the 7,903 bins unique to *xpc-1* XR-seq. 34.7% have been annotated (dark blue) and the remaining 65.3% have not been annotated (light blue) according to the WormBase WS282 annotations. The distribution of chromosomal locations (I-X) is indicated for the unannotated bines. The 68% of annotated bins map to chromosome IV which is not indicated. (C) Pie chart summary of the bins overlapping 2,722 annotations unique to *xpc-1* XR-seq dataset. The majority of the unique annotated bins contain piRNAs from chromosome IV, with the remainder consisting of pseudogenes, protein coding regions, eRNAs, lincRNAs, and nRNAs. The ‘other’ category consists of RNAs excluded from the capped RNA-seq dataset (snRNA, tRNAs, rRNAs) only contains 1.5% of bins. (D) Bin distribution along chromosome IV of unique to the *xpc-1* XR-seq dataset-unannotated bins (top in blue), unique to the *xpc-1* XR-seq dataset-bins with piRNA annotations (middle in burgundy), and intergenic piRNAs from WormBase WS282 annotations (bottom in black).

We further investigated the location and identity of the 7,903 bins that were only detected in the *xpc-1* XR-seq data set and **Fig 6B** shows a pie chart summarizing the results. Of the *xpc-1* XR-seq-unique bins, 34.7% were annotated (dark blue) and the remaining 65.3% have not been annotated (light blue). Of the bins overlapping the 2,722 annotations, 76.8% of those are annotated as piRNAs **(Fig 6C)** and are primarily found on chromosome IV **(Fig 6D)**. Interestingly, 26% of the unannotated bins also map to chromosome IV **(Fig 6C)** and we hypothesize that these may be novel piRNAs or piRNA precursors. In summary, the *xpc-1* XR-seq data set is a useful tool for detecting RNAPII transcription and identifying new transcripts in the previously unannotated intergenic regions of *C. elegans*.

## MATERIALS AND METHODS

### Biological Resources

The *C. elegans* wild-type (N2 ancestral), *csb-1* (RB1801) and *xpc-1* (TG2226) strains were obtained from the *Caenorhabditis* Genetics Center and were cultured under standard conditions at room temperature on nematode growth media (NGM) agar plates with *E. coli* strain OP50.

### XR-seq

To obtain L1 larvae, eggs were collected from adult animals by hypochlorite treatment, and kept in M9 buffer at 22℃ for 16 hours with gentle rotation. Arrested L1 larvae were placed on NGM agar plates with OP50, fed with bacteria for 3 to 4 hours to eliminate the effect of starvation, then exposed to 400 kJ/cm^2^ of UVB radiation (313 nm). The worms were collected in M9 buffer at 5 minutes, 1 hour, 8 hours, 16 hours, 24 hours, and 48 hours after irradiation, and washed until the supernatant became clear. Similarly, mixed-stage worms were exposed to 400kj/cm^2^ of UVB radiation, then collected 1 hour after UVB. The pelleted *C. elegans* (∼50 μl for each) were then incubated for 2 hours at 62°C with 450 μl of Worm Hirt Lysis Buffer (0.15M Tris pH 8.5, 0.1M NaCl, 5mM EDTA, 1% SDS) and 20 μl of Proteinase K (NEB, cat. no. P8107S). Subsequently, 120 μl of 5M NaCl was added, and the mixture was inverted to ensure proper mixing, followed by an overnight incubation and one hour centrifugation at 4°C. Supernatants were processed for XR-seq assay as described previously [20]. In brief, supernatants were incubated with 5μL RNase A and then 5μL Proteinase K, purified, and then immunoprecipitated with either anti-CPD antibody. Immunoprecipitations were ligated to the adaptors, purified with the antibody used in the first purification, and DNA damage was reversed by either CPD photolyase. After PCR amplification, the library was sequenced with either Illumina HiSeq 4000 or NextSeq 2000 platforms.

### RNA-seq

We followed existing protocol [36] for total RNA extracting in *C. elegans*. Briefly, L1 stage wild-type (WT) and *xpc-1 C. elegans* were collected in M9 and washed until the supernatant was clear, followed by incubation with TRizol and chloroform. After centrifugation at 14,000g for 15min at 4°C, the aqueous phase was mixed with an equal volume of isopropanol. Following centrifugation, the RNA pellet was washed several times and then resuspended in RNase-free water. Quality control, followed by stranded and poly(A) enriched library preparation and sequencing, was performed by Novogene.

### Bioinformatic processing

For XR-seq, cutadapt was used to trim reads with adaptor sequence TGGAATTCTCGGGTGCCAAGGAACTCCAGTNNNNNNACGATCTCGTATGCCGTCTTC TGCTTG at the 3′-end and to discard untrimmed reads [37]. Bowtie 2 was used for read alignment to the ce11 reference genome, followed by filtering, sorting, deduplication, and indexing [38]. Post-alignment filtering steps were adopted using Rsamtools (http://bioconductor.org/packages/Rsamtools). We only keep reads that: (i) have mapping quality greater than 20; (ii) are from chromosome I, II, III, IV, V, and X; and (iii) are of length 19-24 bp. Summary statistics of the XR-seq data that we generated are in **S1 Table**. For RNA-seq, reads were aligned using STAR, followed by a filtering step to remove unmapped reads, reads with unmapped mates, reads that do not pass quality controls, reads that are unpaired, and reads that are not properly paired [39]. We only kept the first read from the mate pair to ensure independent measures. Read counts for each gene were obtained using FeatureCounts [40].

### Quality control and data normalization

For gene-specific XR-seq and RNA-seq measurements, we used RPKM for within-sample normalization, since the number of TT dinucleotides are highly correlated with the gene lengths from both the transcribed (TS) and non-transcribed (NTS) strands (**S4 Fig**). To investigate the relationship between gene expression, chromatin states and excision repair, we adopted a stringent quality control (QC) procedure and only retained 26,058 genes that: (i) had at least ten TT dinucleotides in the TS or the NTS; (ii) were less than 300 kb; and (iii) had at least ten reads in total across all XR-seq samples. We observed a robust correlation in repair patterns across the genome between the two replicates collected at each timepoint, underscoring the high reproducibility of our findings (**S11 Fig**). Moreover, pairwise correlation analysis of transcription-coupled repair patterns revealed sample clustering and temporal ordering of samples collected at different time intervals (**S12 Fig**).

To assess excision repair and transcription from non-coding intergenic regions, we generated consecutive and non-overlapping genomic bins of 200 bp long for a total of 501,436 bins. We then removed bins that overlap with annotated genes (gene bodies + 2 kb upstream of the TSS) and those that overlap with blacklist regions in the ce11 genome, resulting in 85,418 bins[41]. For XR-seq, RNA-seq, and short- and long-capped RNA-seq, we adjusted for library size (total number of reads divided by 10^6^) for each bin. When times-series XR-seq data were reported in a combined fashion, we took the median repair across all timepoints to get the CPD repair in replicate 1 and replicate 2, respectively.

### Repair profiles of TS and NTS

For plotting strand-based average repair profiles of the genes in **Fig 2A** and **S3 Fig**, we used WormBase WS282 genome annotations, and filtered 2,142 genes longer than 2 kilobase (kb) pair, situated at least 500 base pairs (bp) away from neighboring genes. For each gene, the region spanning from 1 kb upstream of the TSS to 500 bp downstream was divided into 50 bins. Similarly, the region from 1 kb upstream to 500 bp downstream of the transcription end site (TES) was also divided into 50 bins, resulting in a total of 100 bins per gene. Bed files of the reads were intersected to the 100 bin-divided-gene list by Bedtools intersect with the following commands -c -wa -F 0.5 -S or -s for TS and NTS, respectively^31^. Summary statistics for TCR, measured by TS/(TS+NTS) are represented in **S2 Table**.

To visualize repair around TSS in **Fig 2C** and **S5 Fig**, we filtered 16,588 TSS from WormBase WS291 annotations, which are at least 1 kb apart from each other. We intersected XR-seq reads over 500 bp downstream and upstream of TSS in a strand specific manner. RPKM normalized bigWig files used to create a matrix with the computeMatrix module of deepTools with the following commands reference-point -b 500 -a 500 –missingDataAsZero, and heatmap generated by plotHeatmap module of deepTools [42].

### Identification of dynamic repair using time-course XR-seq data

We next seek to identify genes that exhibit significantly differential and dynamic repair using the time-series XR-seq data of *xpc-1* mutants at 5min, 1h, 8h, 16h, 24h, and 48h in **S6 Fig**. We used Trendy to carry out a breakpoint analysis, allowing for at most two breakpoints and three segments and at least one sample per segment [43]. We used a permutation-based approach with shuffled timepoints to determine the threshold of *R*^2^ (i.e., percentage of total variance that is explained from fitting the time-series model). For the identified significant genes that exhibit dynamic repair across timepoints, we further carried out gene ontology (GO) analysis to identify significantly enriched terms in both biological processes and cellular components [44].

### Capped RNA-seq and epigenomic data

Capped RNA-seq captures nuclear RNAs that are with or without poly(A) tails and is thus much more sensitive in detecting non-coding RNAs compared to RNA-seq. We took advantage of short- and long-capped RNA-seq data of wildtype L1 *C. elegans* that are strand-specific [5]. Additionally, we accessed and cross-compared publicly available epigenomic profiles of L1 *C. elegans*, including chromatin accessibility by ATAC-seq, DNase I hypersensitivity by DNase-seq, and histone modifications (H3K4me1, H3K4me3, and H3K27me3) by ChIP-seq [5]. All data were downloaded as processed bigWig files (**S3 Table**) and lifted over to ce11 when necessary. Regions from the bigWig files were overlapped with the genomic bins, and scores from the bigWig files were averaged, weighted by region widths, to yield the capped RNA-seq and epigenetic measurements for each intergenic region.

### Chromatin state, eRNA, lincRNA, and piRNA annotations

The genic and intergenic regions of *C. elegans* (ce11) were annotated using the GenomicFeatures R package in conjunction with the TxDb.Celegans.UCSC.ce11.refGene annotation package. Chromatin states in the L3 stage of *C. elegans* were previously inferred, consisting of 20 distinct states as detailed in **Fig 4A** and **S7 Fig** [45]. Evans *et al*. observed a high degree of similarity in autosomal chromatin states between the embryonic and L3 larval stages of the worms. This conservation of chromatin configuration allowed us to confidently use the chromatin state data from the L3 stage for intersection with our L1 stage data, without compromising the integrity of our analysis [45]. Each annotated chromatin region was mapped from ce10 to ce11 and intersected with RNA-seq, capped RNA-seq, and XR-seq reads. For eRNAs, 90 % of which are bidirectionally transcribed, non-polyadenylated and unspliced, we retrieved 505 annotated eRNAs in *C. elegans* from the eRNAdb database [46, 47]. We removed eRNAs that overlap with either annotated genes or blacklist regions, resulting in a total of 324 eRNAs, which are presented in **Fig 5A** and **S9A Fig**. Similarly, we obtained 170 long intergenic non-coding RNAs (lincRNAs) in *C. elegans* from existing annotations [48]. After lifting over the coordinates from ce6 to ce11 and filtering out ones that overlap with genes or blacklist regions, we were left with 103 lincRNAs, which are visualized in the **Fig 5B** and **S9B Fig**. We obtained 15,363 piRNAs in *C. elegans* from existing WormBase WS282 annotations. Removing the piRNAs that overlap with genes or blacklist regions results in 10,757 intergenic piRNAs, which are shown in **Fig 5C** and **S9C Fig**.

## DISCUSSION

Transcription-coupled repair appears to be universal in cellular organisms ranging from bacteria to humans and has been studied in several model organisms [10, 22, 24, 49–54]. Multiple methodologies have been developed to unravel the intricate mechanisms and required repair factors [13]. Among these methods, XR-seq, distinguished by its whole-genome analysis at single-nucleotide resolution, has been applied across a spectrum of organisms, including bacteria, yeast, flies, plants, and mammals [13]. A previous study employing a qPCR assay, indicated the existence of transcription-coupled repair in *C. elegans* [16], nevertheless, our study stands as a single-nucleotide-resolution genome-wide UV-damage transcription-coupled repair map of this important model organism. Furthermore, our investigation distinguishes itself by employing transcription-coupled repair as a proxy for RNAPII transcription, and *xpc-1* XR-seq data to effectively complement RNA-seq and capped RNA-seq datasets to offer a more comprehensive view of transcription.

Leveraging the unique properties of XR-seq data, we aimed to delve into the realm of intergenic transcription, a domain that has posed persistent challenges for conventional RNA-seq methods. Based on the RNAPII disassociation model in response to UV-induced damage, RNAPII encounters transcription blockage and initiates a process of transcription-coupled repair. During this repair process, RNAPII dissociates from the DNA strand, facilitating the sequential removal of lesions from the template in the 5’ to 3’ direction. This concerted repair mechanism eventually leads to the clearance of adducts from the template, thereby enabling the synthesis of full-length transcripts [26, 55]. To comprehensively investigate these intricate transcription dynamics, we conducted XR-seq at six distinct timepoints, ranging from 5 minutes to 48 hours following UV treatment. As a result, our dataset encompasses both transcription initiation and elongation events, providing a comprehensive view of the entire transcriptional process.

Detection of non-coding RNAs has long been a formidable task due to their relatively low abundance and inherent instability. The development of cutting-edge technologies, such as RNA polymerase II chromatin immunoprecipitation coupled with high-throughput sequencing (RNAPII ChIP-seq), Global Run-On sequencing (GRO-seq), Precision Run-On Sequencing (PRO-seq), and a variety of methods for sequencing the 5’-anchored RNAs, has been driven by the desire to discern nascent RNAs and ncRNAs with heightened precision [5, 6, 18, 56–59]. A comprehensive evaluation of the strengths and limitations of these methods has been described elsewhere [60], and in the context of *C. elegans* research, efforts to specifically target ncRNAs and identify TSS have utilized 5’-capped RNA-sequencing methods, as reported in previous studies [6, 18, 45, 61–63].

XR-seq presents a noteworthy advantage in its ability to directly detect transcription events at the DNA level, thus circumventing the inherent limitations associated with indirect transcription detection techniques such as RNA sequencing. These conventional methods are prone to challenges stemming from the low abundance and instability of RNA molecules. Furthermore, RNA sequencing is susceptible to sequence bias resulting from early transcriptional events that introduce differences between RNA and DNA sequences [64, 65]. XR-seq, conversely, by its nature of sequencing transcribed DNA, effectively eliminates this sequence bias, ensuring a more accurate representation of transcriptional activity. An additional advantage of XR-seq is its applicability to prokaryotic organisms, mirroring its utility in eukaryotes, a distinction not shared by nascent RNA sequencing methods.

Our findings demonstrate the efficacy of XR-seq in capturing transcription events within both genic and intergenic regions. While RNA-seq detects only 17.5% of intergenic transcription, our data reveal that up to 80% of the overall intergenic transcription landscape is covered and shared between XR-seq and capped RNA-seq. Notably, XR-seq exhibits sensitivity comparable to that of capped RNA-seq in detecting annotated intergenic enhancer RNAs (eRNAs) and long intergenic non-coding RNAs (lincRNAs), but is superior at detecting intergenic Piwi-interacting RNAs (piRNAs). In *C. elegans*, piRNAs are transcribed from >15,000 discrete genomic loci by RNAPII, resulting in 28-nt short-capped piRNA precursors that play key roles in germline development, genome integrity, and other biological processes [66–69]. The majority of piRNAs are localized to two ∼3 Mb cluster regions on chromosome IV [70], and we found 2,090 annotated piRNAs and 1,341 unannotated intergenic regions unique to *xpc-1* XR-seq on this chromosome. We hypothesize that many of these unannotated intergenic regions on chromosome may either be transient piRNA precursor transcripts not captured by other methods or that they are UV-induced piRNAs. Future studies using methods that efficiently capture piRNAs, such as CIP-TAP [18], CAGE [70] or short capRNA-seq [71], after exposing worms to UV could be very informative. In conclusion, our findings provide valuable insights into nascent transcription dynamics and the intricate interplay between transcription-coupled repair and intergenic transcription in *C. elegans*, and this knowledge will be valuable when translated to the human genome and other organisms with large unmapped intergenic content.

## AUTHOR CONTRIBUTIONS

A.S. and L.L.-B. envisioned and initiated the study, while C.K. conducted the experiment. All authors designed and conducted the analysis, wrote, and approved the manuscript.

## DATA AVAILABILITY

XR-seq and RNA-seq data reported in this paper have been deposited in the Gene Expression Omnibus (GEO) database with accession number GSE245181 and GSE262486. ATAC-seq, ChIP-seq, and DNase-seq are available from GEO with accession numbers GSE114439, GSE114440, and GSE114481, respectively. All code used in this paper is available at https://github.com/yuchaojiang/damage_repair/tree/master/C_elegans.

## ACKNOWLEDGEMENTS

This work was supported by National Institute of health grants R35 GM118102 (A.S.), R01 ES033414 (A.S.) and R35 GM138342 (Y.J.). Portions of this research were conducted with the advanced computing resources provided by Texas A&M High Performance Research Computing. The authors thank Dr. Shawn Ahmed for helpful discussions and comments.

