## Supplement for "Genome-wide analysis of transcription-coupled repair reveals novel transcription events in *Caenorhabditis elegans*"

**S1 Fig. The Excision Repair-sequencing (XR-seq) Method.** Excised oligonucleotides isolated by Hirt lysis from lysed worms, purified with anti-CPD specific antibodies, ligated to adapters, and then purified with anti-CPD specific antibodies to remove excess adaptors. The damage was then reversed with CPD photolyase and then PCR was performed to generate libraries for high throughput sequencing.

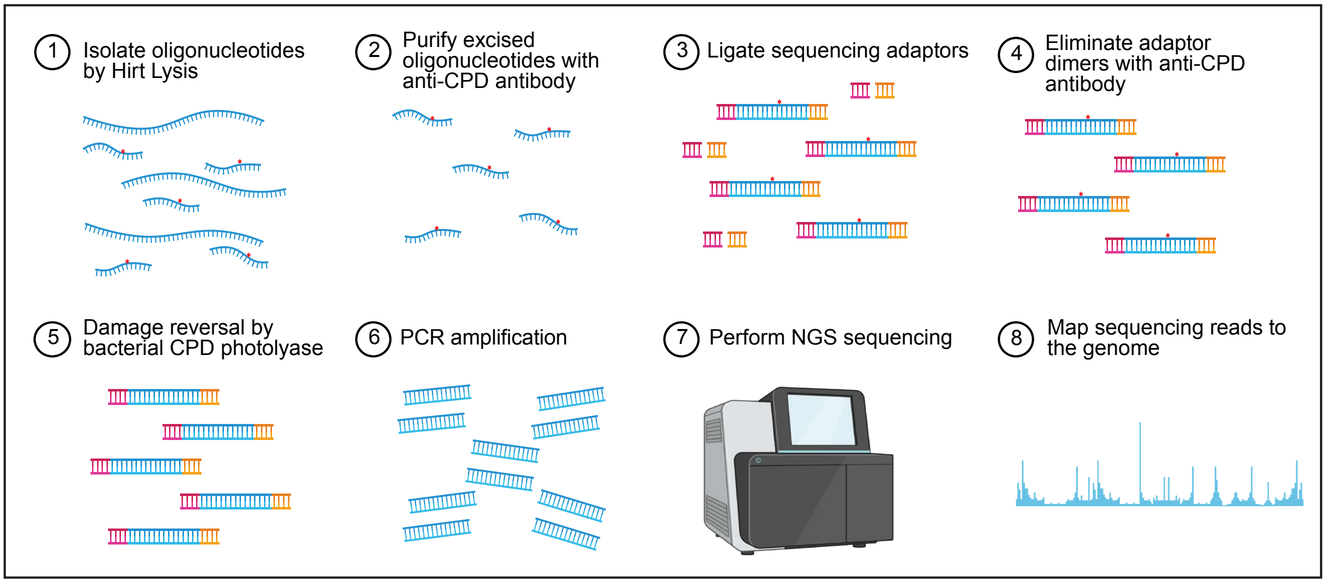

**S2 Fig. Nucleotide distribution in XR-seq reads of 19-30 nt from wild-type, *csb-1,* and *xpc-1* at 1 hour repair time.** The enrichment of TT is observed at a fixed distance, 6 nt from 3' end, indicating that CPD-damage carrying oligonucleotides are successfully represented by XR-seq.

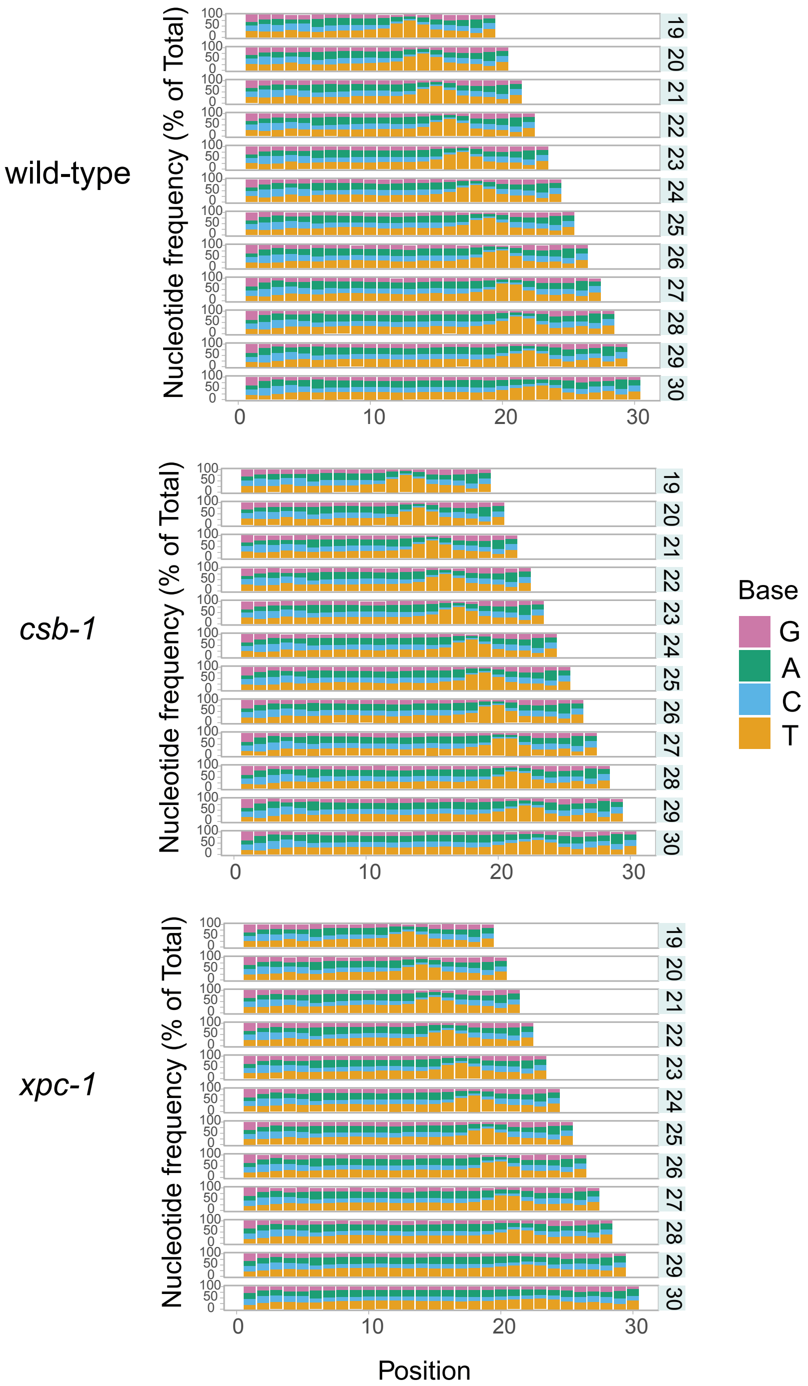

**S3 Fig. Nucleotide distribution and genome-wide repair of transcribed strand (TS) and non-transcribed strands (NTS) in *xpc-1* XR-seq. (A)** Time-course of *xpc-1* XR-seq, shows TT enrichment 6 nt from 3' end in reads of 24 nt. **(B)** Genome-wide transcribed and non-transcribed strand repair in time-course *xpc-1* XR-seq is plotted as average RPKM (y-axis) along the x-axis 500 bp upstream and 1 kb downstream of transcription start sites (TSS), and 1 kb upstream and 500 bp downstream of transcription end site (TES) for 2,142 genes selected for length > 2 kb and no overlaps with a distance of at least 500 bp between genes. **(C)** XR-seq from *xpc-1* mixed stage worms 1h after UV showing nucleotide distribution and genome-wide transcribed and non-transcribed strand repair as in A and B above.

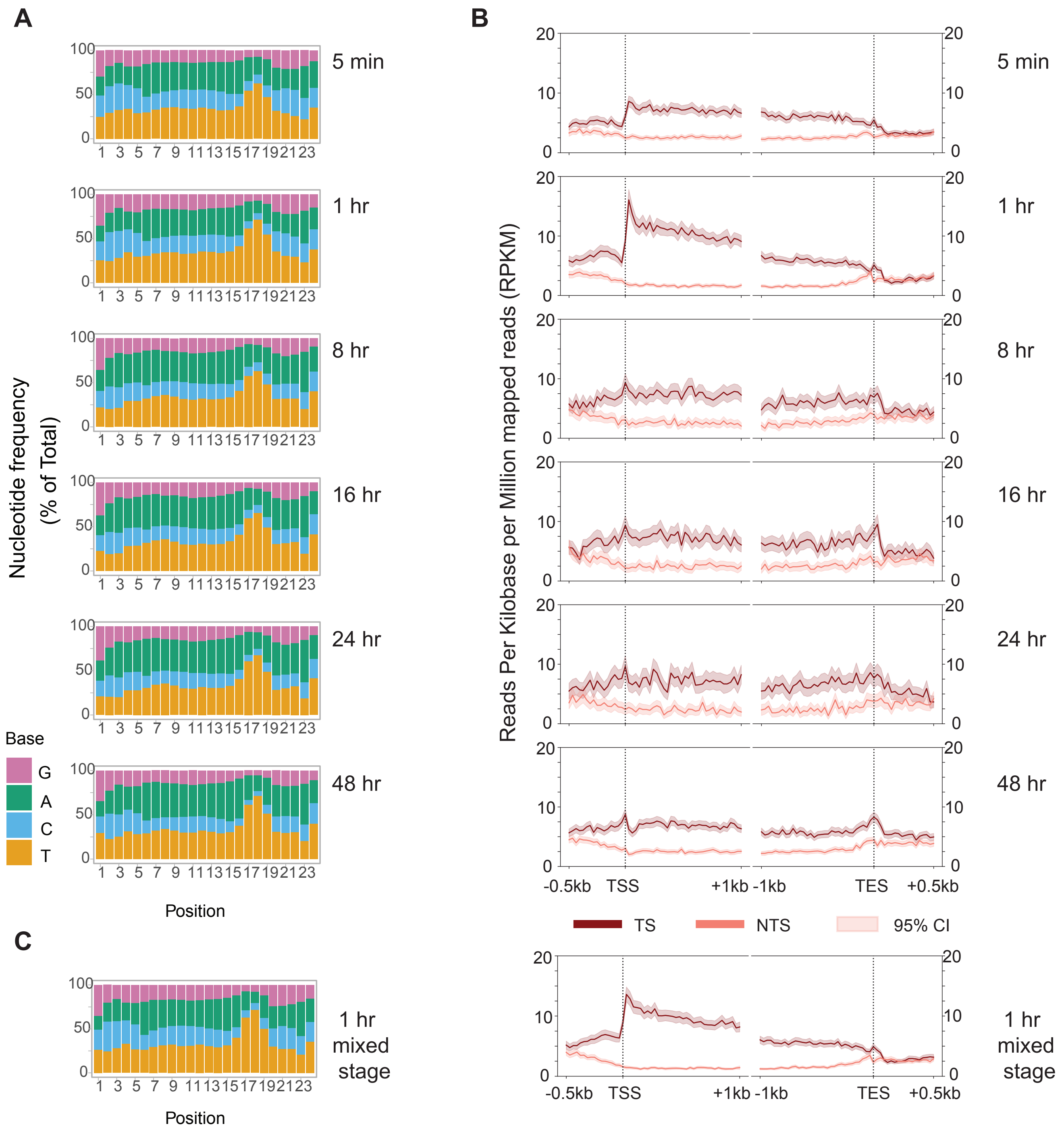

**S4 Fig. TT dinucleotides in the transcribed strand (TS) and non-transcribed strand (NTS).** The number of TT dinucleotides in the TS is highly correlated with that in the NTS. Natural logarithms of the numbers of TT dinucleotides from the TS and NTS were computed, respectively. Gene length is a good proxy for the number of TT dinucleotides, and thus we use RPKM for normalization for XR-seq repair read counts.

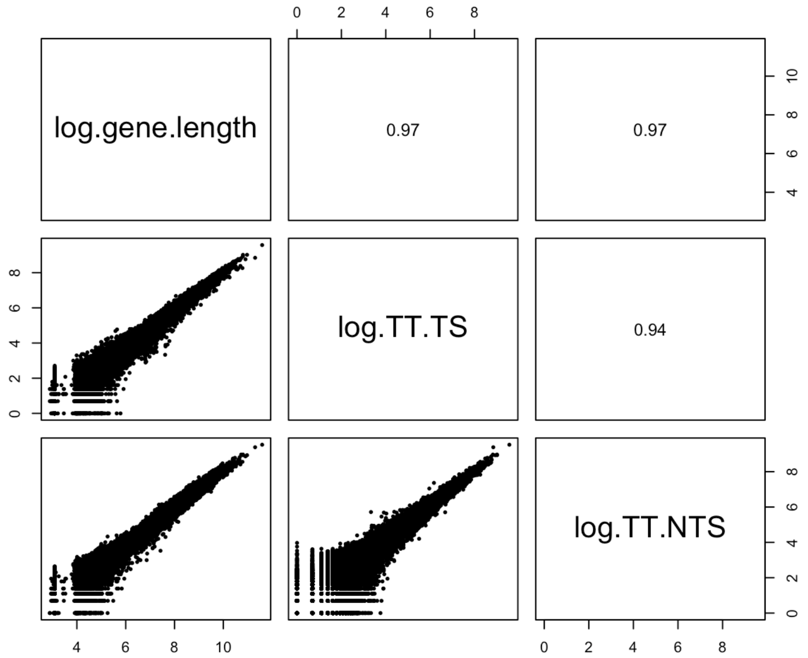

**S5 Fig. Extended Figure 2C showing repair in the TS and NTS around TSSs in the wild-type (L1), *csb-1* (L1) and *xpc-1* (L1 and mixed) XR-seq, 1 h after UV.** In wild-type, upstream of TSS has more repair on NTS, in contrast to more repair in TS downstream of TSS. In *csb-1,* despite more repair on the NTS upstream of TSS, TSS downstream repair does not show a strand preference. In *xpc-1*, repair in L1 worms and mixed stage worms exhibit similar profiles, proving that the repair preference in TS at TSS and its immediate downstream is not unique to the L1 worms. Near background repair at NTS versus efficient repair at TS is additional evidence of lacking global repair in *xpc-1*. Although profile plots (top) mask the anti-sense transcription-coupled repair upstream of TSS, a subset of TSSs exhibits upstream TCR on the non-template strand. Genome-wide TT content (right) across the same selected TSSs shows a dip in both strands at TSS. There are more TT dinucleotides on the NTS than TS upstream of TSSs, and therefore more theoretical damages which result in more repair reads in wild-type and *csb-1* in that region.

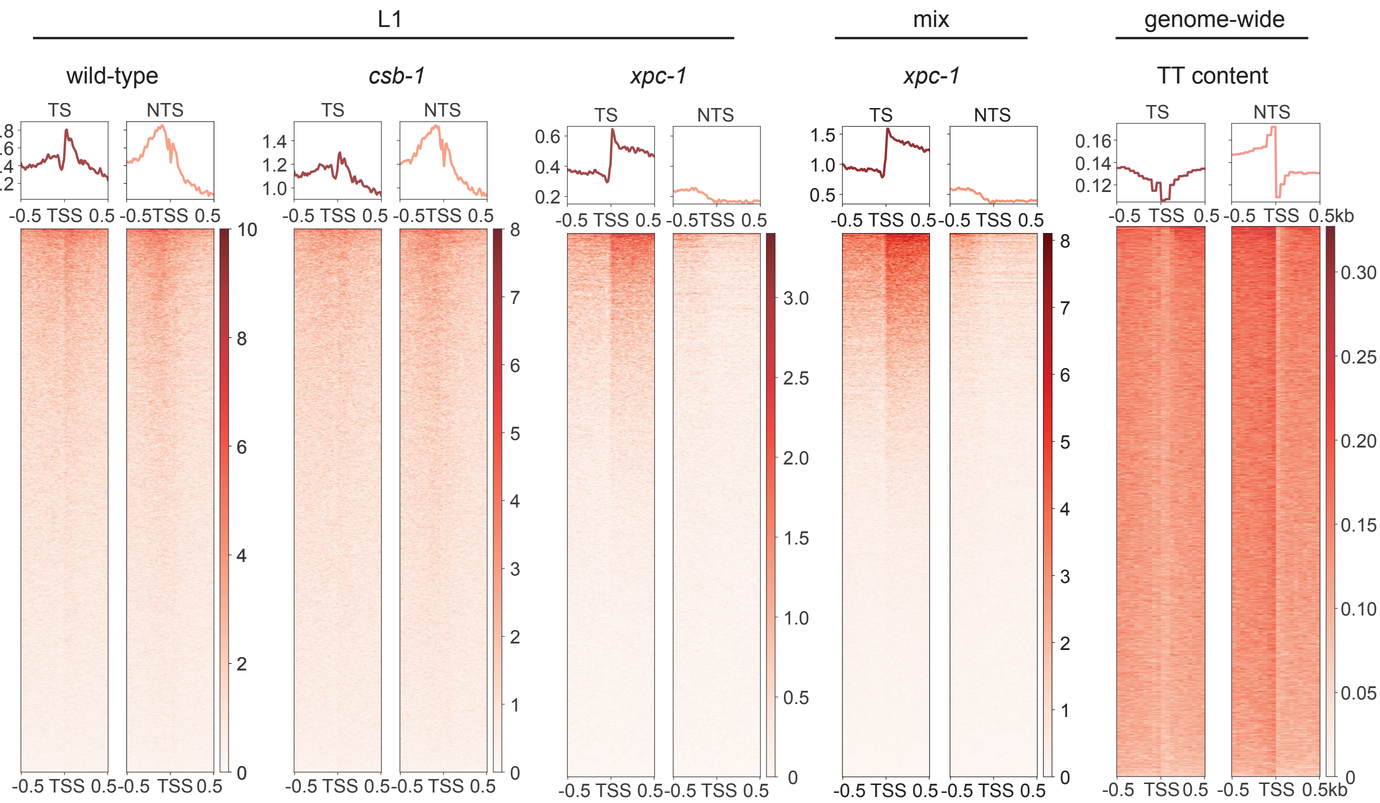

**S6 Fig. Time-course differential-repaired genes.** Two XR-seq replicates of *xpc-1* collected at 5min, 1h, 8h, 16h, 24h, and 48h were included in the breakpoint analysis. **(A)** Heatmap of 121 significant genes that show dynamic repair patterns across timepoints. Genes cluster into two clades that exhibit early (112 genes) and late repair (9 genes). Early-repair genes were tested for gene-ontology enrichment, with significantly enriched terms shown in **(B)** biological process and **(C)** cellular component.

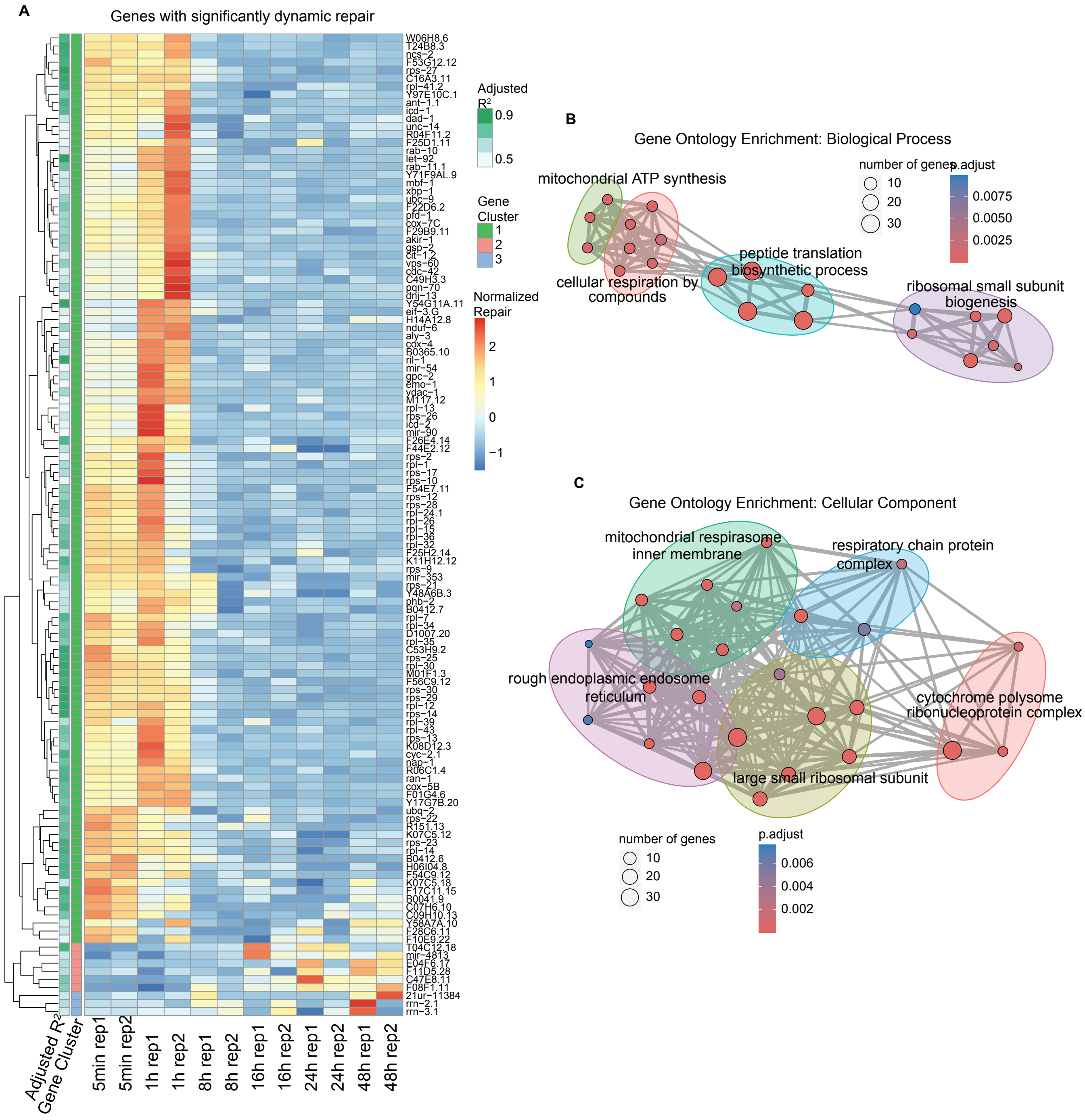

**S7 Fig. Genome-wide distribution and epigenetic signatures of the reads from RNA-seq, capped RNA-seq, and XR-seq. (A)** Extended Figure 3B bar graphs depicting the genome-wide distribution of reads obtained from various sequencing methods, including wild-type (WT) and *xpc-1* RNA-seq, long-capped RNA-seq, short-capped RNA-seq, and WT, *csb-1, xpc-1* XR-seq. Notably, both XR-seq and capped RNA-seq techniques reveal transcription events occurring outside of genic regions. **(B)** Extended Figure 4A heatmap with reads from wild-type (WT) and *xpc-1* RNA-seq, long-capped RNA-seq, short-capped RNA-seq, and WT, *csb-1, xpc-1* XR-seq were overlapped with genomic intervals corresponding to 20 distinct chromatin states predicted for the autosomes of *C. elegans*. Proportion of reads were computed for each of the annotated chromatin states; square root of the proportion is visualized as a heatmap.

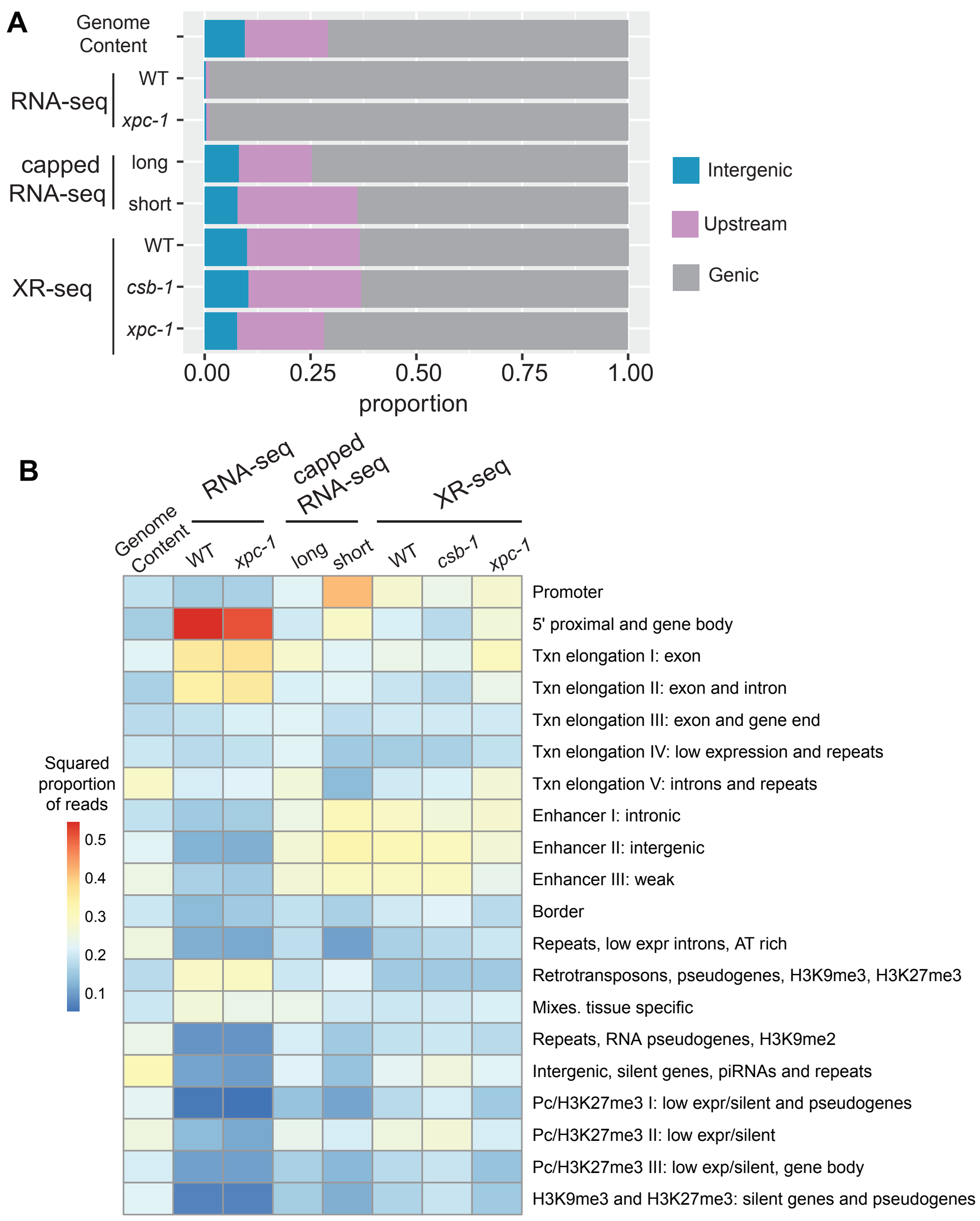

**S8 Fig. Repair events captured by XR-seq are highly correlated with the capped RNA-seq transcription events.** XR-seq repair signals correlate with short- and long-capped RNA-seq signals much stronger than conventional RNA-seq. Pairwise smooth scatterplots are shown on the lower triangle, where color corresponds to smoothed data density; Spearman correlation coefficients are shown on the upper triangle, with text size proportionate to the absolute value of the coefficient. Library-size-adjusted read counts from the filtered genomic bins are plotted on the original scale; XR-seq replicates were merged by taking the average, and the 1h timepoint for *xpc-1* was used.

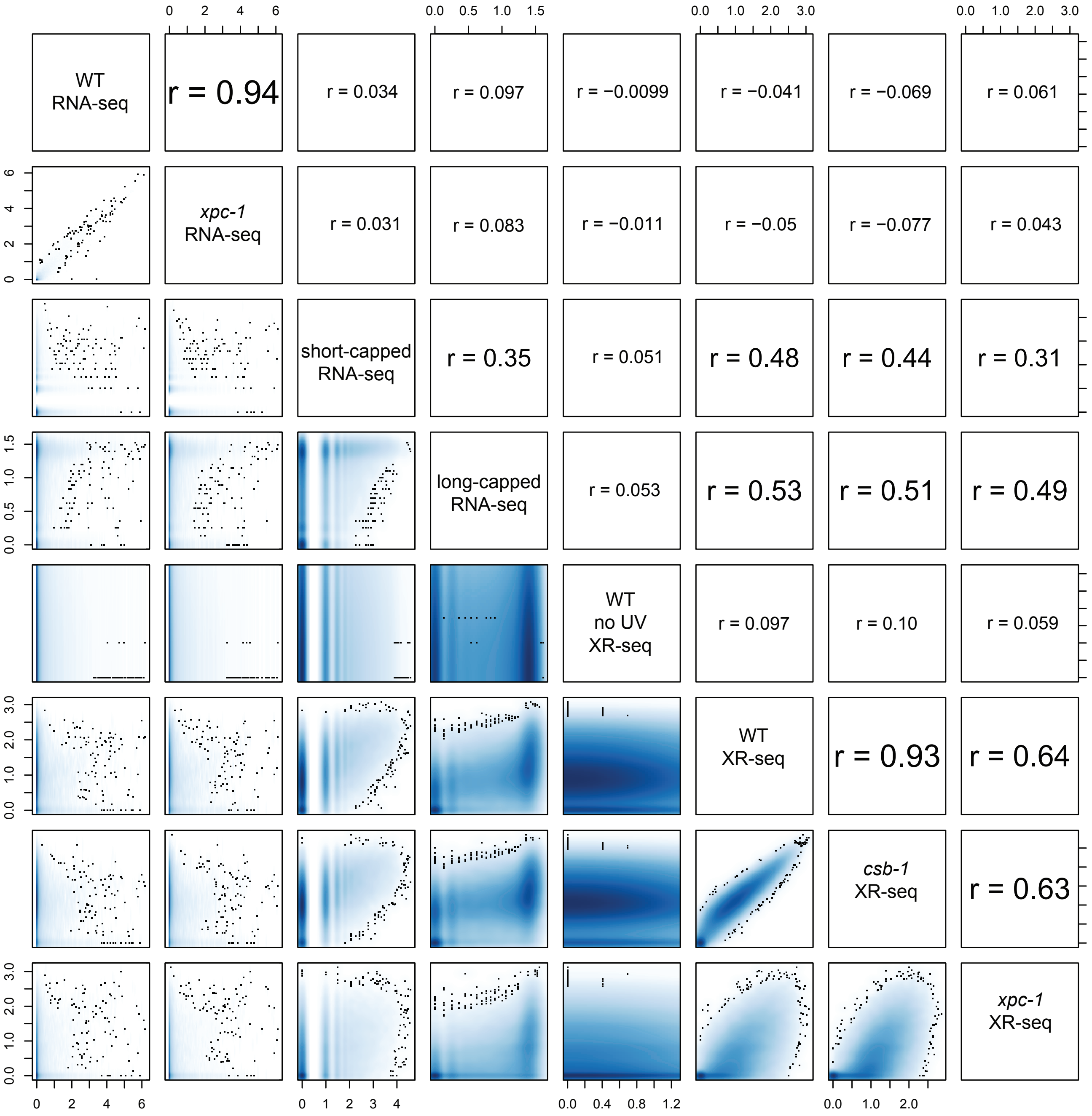

**S9 Fig. Extended Figure 5 analysis of read numbers in intergenic eRNA, linc RNA, and intergenic piRNAs.** **(A)** Heatmaps (left) display normalized reads for intergenic enhancer RNAs (eRNAs) segregated by chromosomes. Normalization by log(x+1) was carried out, where x is library-size-adjusted read count. Bar graphs (right) represent log-normalized read counts for eRNA. Data are presented for WT and *xpc-1* RNA-seq, WT long- and short-capped RNA-seq, and 2 replicates each of XR-seq from WT no UV, 1 hour after UV in WT and *csb-1,* and *xpc-1* combined time-course (5min, 1h, 8h, 16h, 24h, and 48h). **(B, C)** Heatmaps and bar graphs as in A, for long intergenic non-coding RNAs (lincRNAs) and intergenic Piwi-interacting RNAs (piRNAs), respectively.

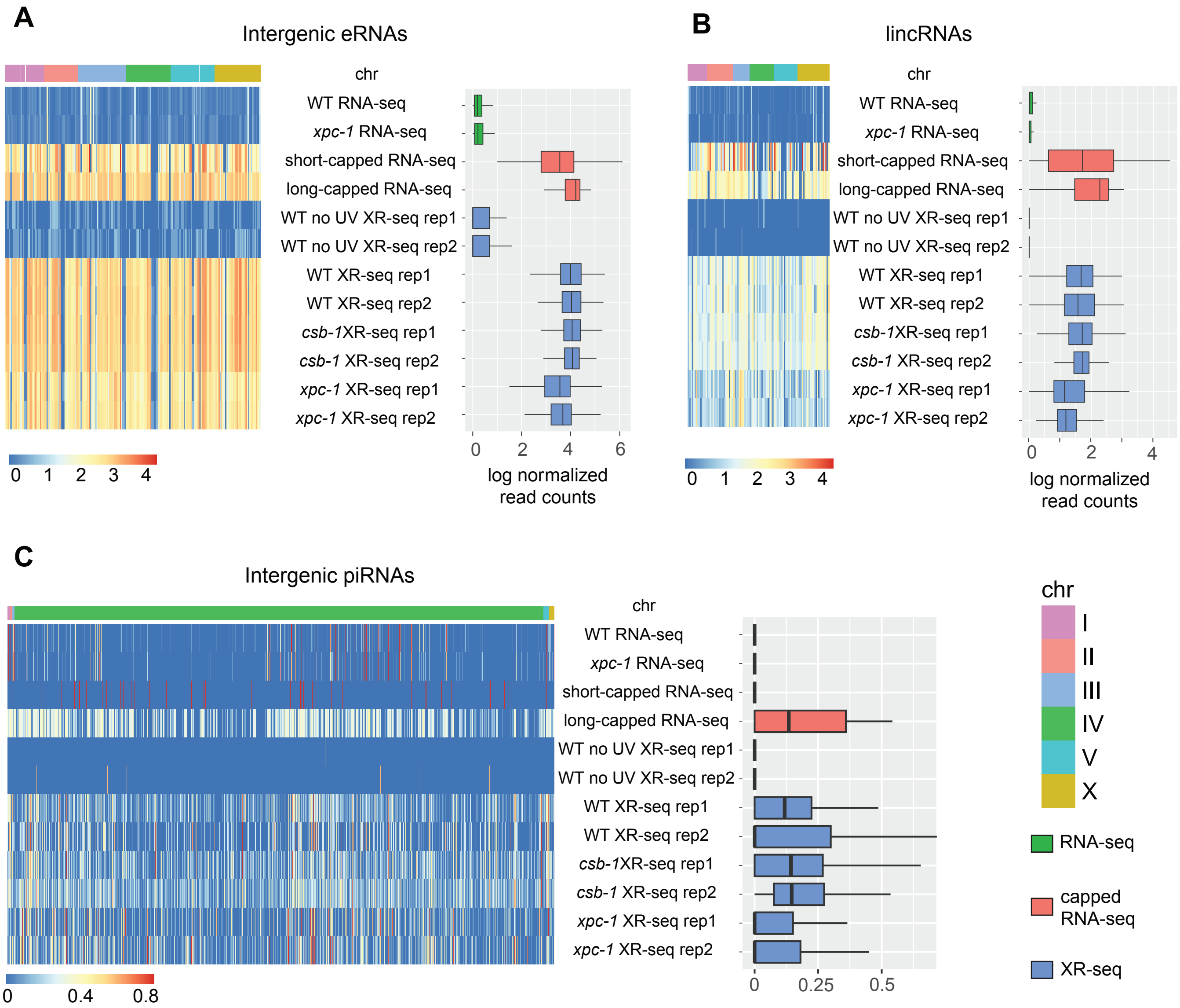

**S10 Fig. Extended Figure 6A showing intergenic repair in the wild-type (L1), *csb-1* (L1) and *xpc-1* (L1 and mixed) XR-seq, 1 h after UV.** For the 85,418 intergenic bins, we identified regions with non-zero read counts by short- or long-capped RNA-seq, RNA-seq, and CPD XR-seq, respectively. **(A)** Upset plot to show the intergenic bins detected by capped RNA-seq, conventional RNA-seq, and XR-seq. To reduce the number of call sets, we required non-zero read counts to be detected: (i) in both replicates for XR-seq; (ii) in both WT and *xpc-1* RNA-seq, as they are highly correlated; and (iii) by either short-capped or long-capped RNA-seq, as they are complementary. **(B)** We used experimental results from short- and long-capped RNA-seq as ground truths and calculated sensitivity, specificity, and F measure (geometric mean of sensitivity and specificity as a joint metric) for the other data types and genotypes. RNA-seq has the lowest sensitivity; *csb-1* XR-seq has the highest sensitivity due to the pervasive global repair detected from intergenic regions, although it also suffers from low specificality.

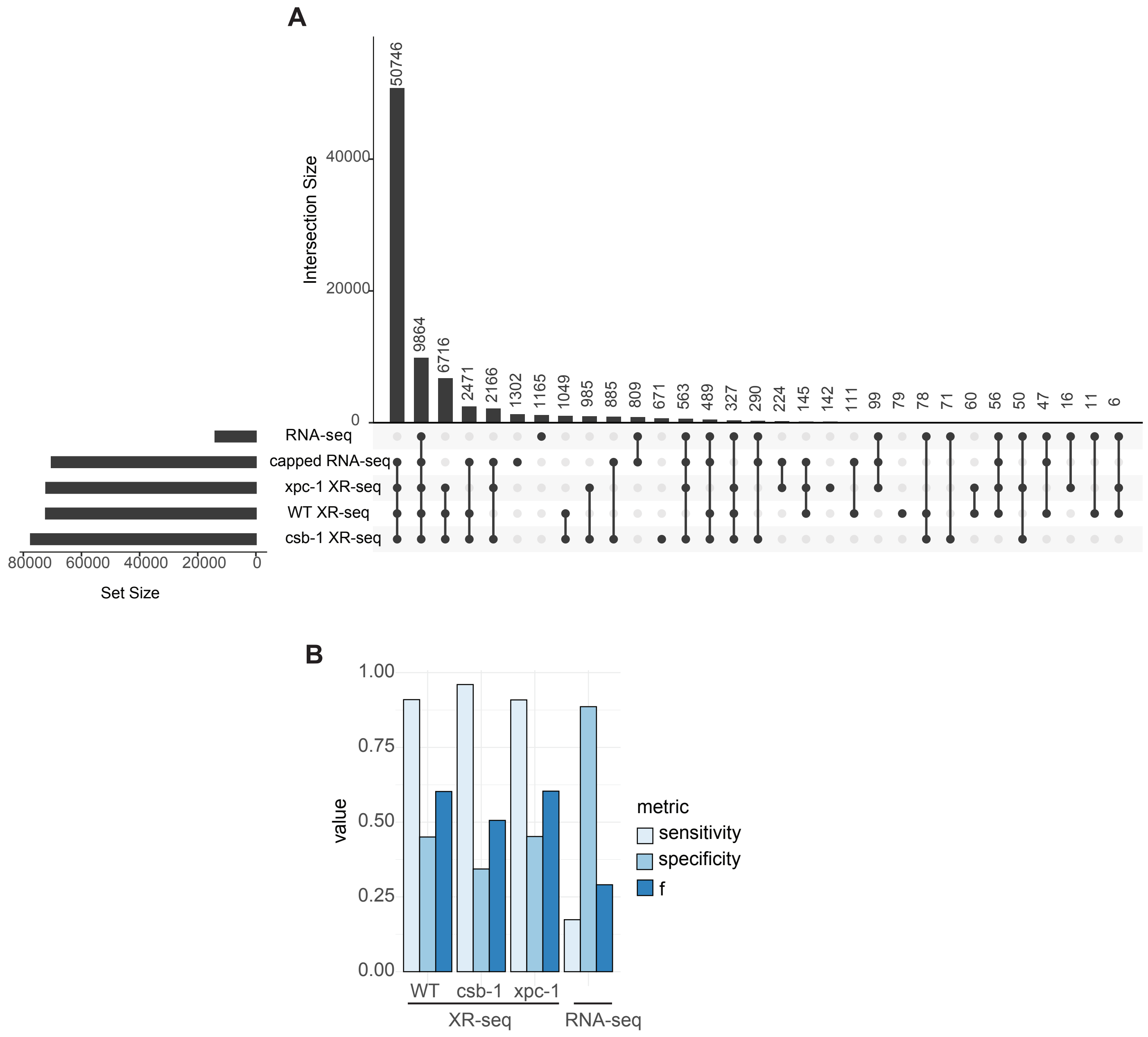

**S11 Fig. High reproducibility between each pair of XR-seq replicates.** Normalized gene-specific repair is shown as each dot. Spearman correlation coefficient is shown.

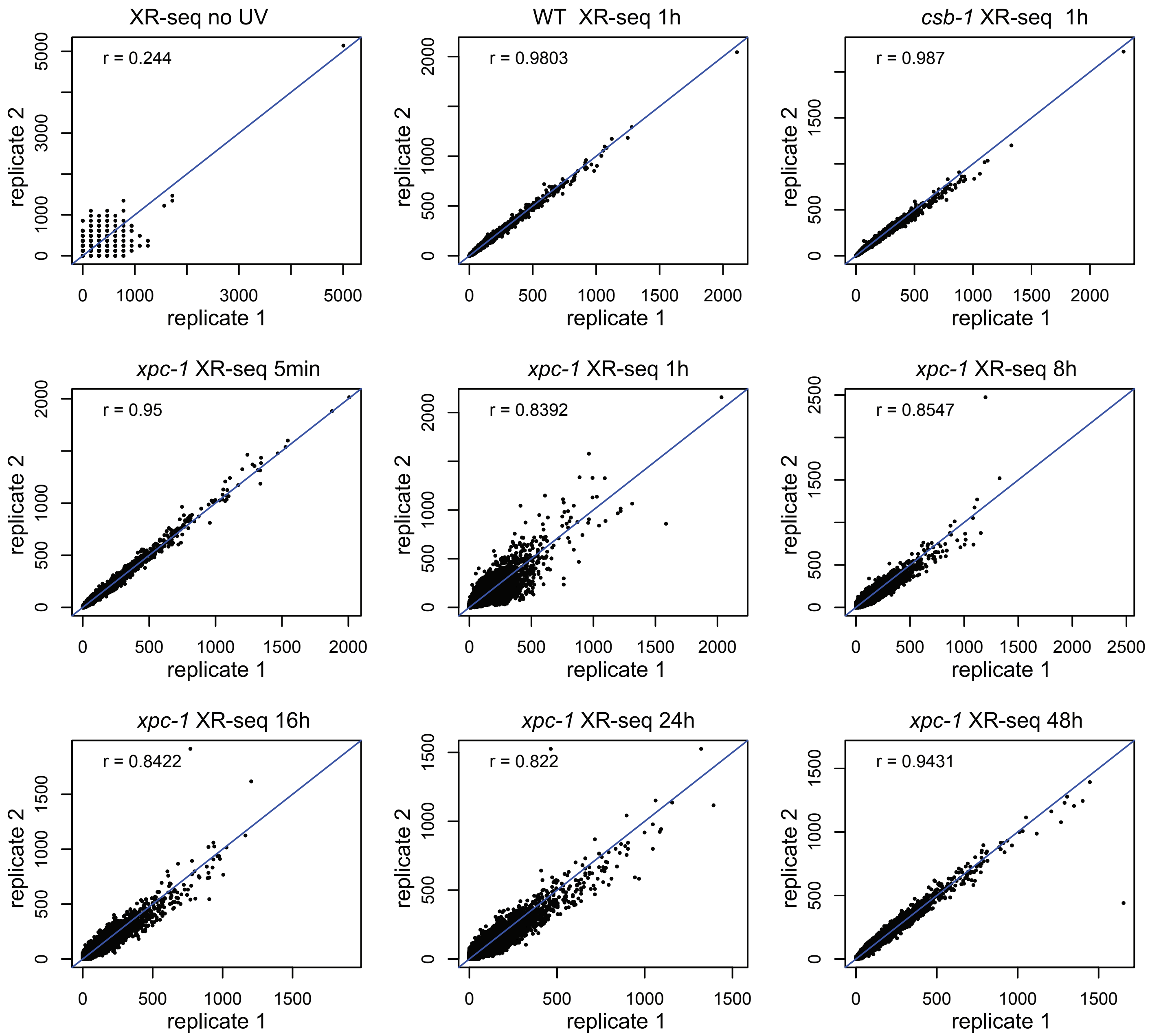

**S12 Fig. Pairwise correlation and principal component analysis of XR-seq reads. (A)** Spearman correlation coefficient is calculated between each pair of the XR-seq samples using the normalized read counts. Heatmap is generated to visualize the symmetric matrix of correlation coefficient. **(B)** First two principal components from the principal component analysis. Each set of repeats clustered closely, and the WT and *csb-1* sets clustered closer to each other than either *xpc-1* or no UV. Between different *xpc-1* timepoints temporal changes were observed.

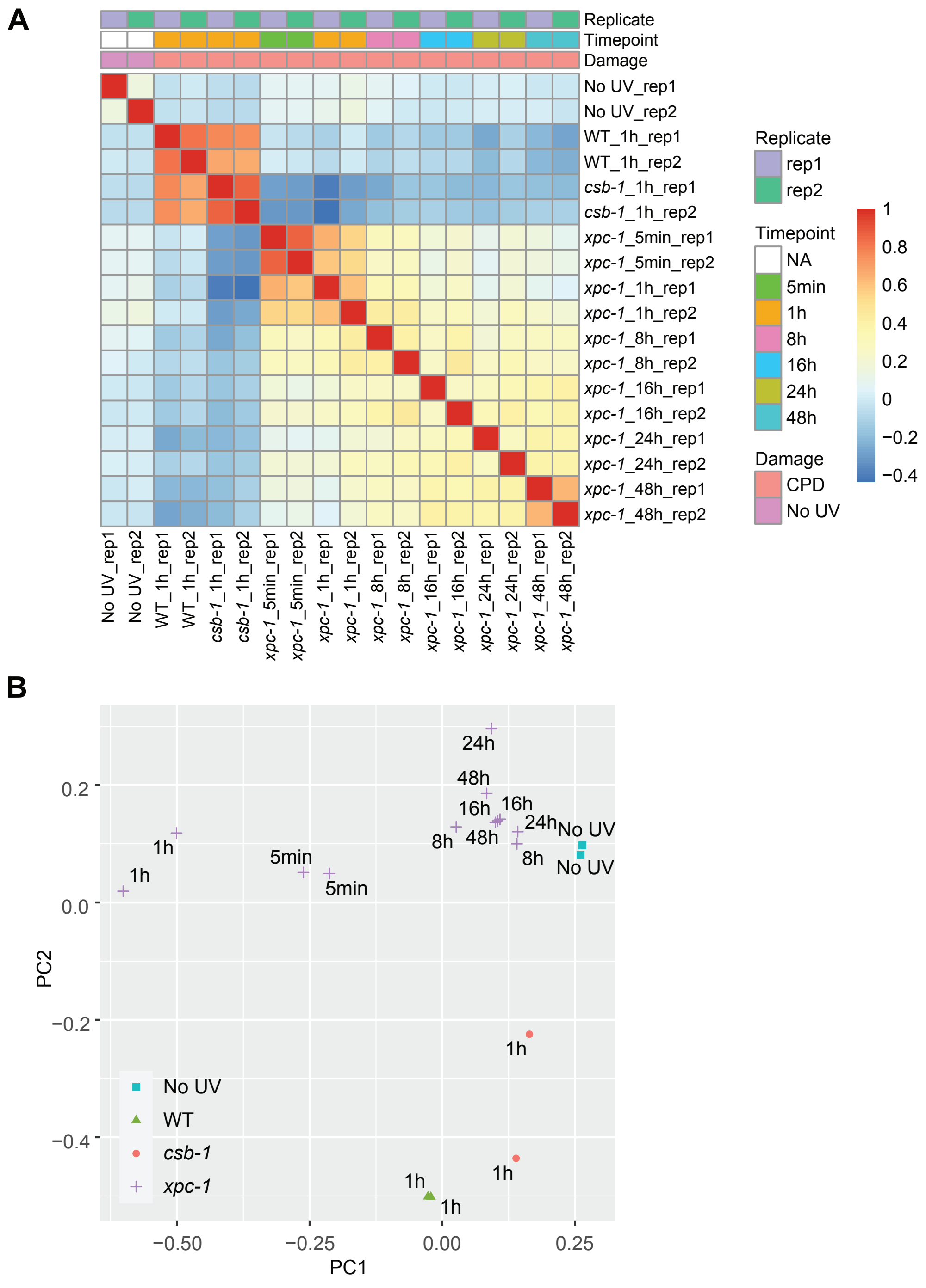

**S1 Table. XR-seq sample information.** Summary of C. elegans (6-4)PP and CPD XR-seq samples across different timepoints and replicates. Total_mapped: total mapped reads. Dedup: deduplicated reads. Mapq: reads with mapping quality > 20 (reads that are equally mapped to multiple genomic locations are removed with this QC). Chr: reads mapped to chrI, II, III, IV, V, X. Qwidth: reads with lengths 19-24. GenebodyPromoter: reads mapped to genes and 2 Kb upstream of transcription start sites (i.e., promoters). Genebody: reads mapped to genes.

| Sample | Genotype | Damage | Timepoint | Replicate | UVB | Total_mapped | Dedup | Mapq | Chr | Qwidth | Genebodypromoter | Genebody |
|---|---|---|---|---|---|---|---|---|---|---|---|---|
| XR_seq_WT_noUV_C_elegans_CPD_rep1 | WT | No_UV | NA | rep1 | 0 | 459991 | 458110 | 24411 | 24334 | 10144 | 8820 | 6244 |
| XR_seq_WT_noUV_C_elegans_CPD_rep2 | | | | rep2 | 0 | 450109 | 447722 | 28609 | 28503 | 12785 | 11114 | 7969 |
| XR_seq_WT_C_elegans_CPD_1h_rep1 | | CPD | 1h | rep1 | 4000 | 21236673 | 20736814 | 16259281 | 16243756 | 7971377 | 6861612 | 4429060 |
| XR_seq_WT_C_elegans_CPD_1h_rep2 | | | | rep2 | 4000 | 8936703 | 8165309 | 5939887 | 5936339 | 2862800 | 2467345 | 1624521 |
| XR_seq_CSB_C_elegans_CPD_1h_rep1 | CSB | CPD | 1h | rep1 | 4000 | 18325556 | 18040700 | 13791578 | 13783275 | 6501325 | 5551605 | 3547344 |
| XR_seq_CSB_C_elegans_CPD_1h_rep2 | | | | rep2 | 4000 | 37970895 | 36961311 | 27920512 | 27899498 | 12726720 | 10871991 | 7050716 |
| XR_seq_XPC_C_elegans_CPD_5min_rep1 | XPC | CPD | 5min | rep1 | 4000 | 5837318 | 5763808 | 4072843 | 4053858 | 1452895 | 1300422 | 981492 |
| XR_seq_XPC_C_elegans_CPD_5min_rep2 | | | | rep2 | 4000 | 4302562 | 4264267 | 2958176 | 2944500 | 1067358 | 954165 | 719738 |
| XR_seq_XPC_C_elegans_CPD_1h_rep1 | | | 1h | rep1 | 4000 | 1717268 | 1710614 | 524924 | 522437 | 227281 | 211003 | 166995 |
| XR_seq_XPC_C_elegans_CPD_1h_rep2 | | | | rep2 | 4000 | 5843021 | 5804098 | 4092344 | 4077700 | 1921303 | 1731004 | 1274284 |
| XR_seq_XPC_C_elegans_CPD_8h_rep1 | | | 8h | rep1 | 4000 | 1340986 | 1327771 | 746362 | 743063 | 314802 | 280403 | 201968 |
| XR_seq_XPC_C_elegans_CPD_8h_rep2 | | | | rep2 | 4000 | 3094513 | 3042372 | 1355727 | 1346939 | 403994 | 355247 | 250147 |
| XR_seq_XPC_C_elegans_CPD_16h_rep1 | | | 16h | rep1 | 4000 | 1082126 | 1072088 | 590778 | 588731 | 259173 | 230254 | 165860 |
| XR_seq_XPC_C_elegans_CPD_16h_rep2 | | | | rep2 | 4000 | 1704743 | 1656693 | 982544 | 976155 | 357955 | 315907 | 224116 |
| XR_seq_XPC_C_elegans_CPD_24h_rep1 | | | 24h | rep1 | 4000 | 684978 | 679498 | 419344 | 417964 | 196339 | 173972 | 123874 |
| XR_seq_XPC_C_elegans_CPD_24h_rep2 | | | | rep2 | 4000 | 1404716 | 1389973 | 814922 | 810876 | 316865 | 278957 | 197723 |
| XR_seq_XPC_C_elegans_CPD_48h_rep1 | | | 48h | rep1 | 4000 | 2943738 | 2844139 | 1969532 | 1952041 | 1023631 | 897604 | 619118 |
| XR_seq_XPC_C_elegans_CPD_48h_rep2 | | | | rep2 | 4000 | 5145059 | 5055556 | 3671400 | 3654971 | 1685725 | 1484541 | 1041861 |

**S2 Table. Transcription-coupled repair (TCR) measured by XR-seq.** The ratio of read counts from the TS to those from both the TS and NTS serves as a proxy for TCR. XPC mutants exemplified the strongest TCR, while CSB mutants showed depleted TCR as expected. TCR in WT samples was mixed with global repair, while TCR in WT samples without UV treatment reflected background noise.

| Genotype | Damage | Timepoint | Replicate | Transcription-Coupled Repair: TS/(TS+NTS) | | |
|---|---|---|---|---|---|---|
| | | | | Lower.Quartile | Median | Upper.Quartile |
| WT | No_UV | NA | rep1 | 0 | 0.5 | 1 |
| | | | rep2 | 0 | 0.5 | 1 |
| | CPD | 1h | rep1 | 0.5083 | 0.5517 | 0.6072 |
| | | | rep2 | 0.5055 | 0.5619 | 0.6269 |
| CSB | CPD | 1h | rep1 | 0.4651 | 0.501 | 0.5442 |
| | | | rep2 | 0.4735 | 0.5035 | 0.536 |
| XPC | CPD | 5min | rep1 | 0.6416 | 0.7802 | 0.859 |
| | | | rep2 | 0.6341 | 0.7808 | 0.8648 |
| | | 1h | rep1 | 0.7143 | 0.875 | 0.9697 |
| | | | rep2 | 0.7165 | 0.831 | 0.9066 |
| | | 8h | rep1 | 0.5833 | 0.75 | 0.8571 |
| | | | rep2 | 0.5294 | 0.6667 | 0.7826 |
| | | 16h | rep1 | 0.6209 | 0.754 | 0.8667 |
| | | | rep2 | 0.5667 | 0.7059 | 0.8235 |
| | | 24h | rep1 | 0.6219 | 0.7826 | 0.9 |
| | | | rep2 | 0.5714 | 0.7044 | 0.8122 |
| | | 48h | rep1 | 0.5714 | 0.6923 | 0.7954 |
| | | | rep2 | 0.5985 | 0.7244 | 0.8243 |

**S3 Table. Epigenomic and capped RNA-seq data of L1 C. elegans adopted in this study.**

| **Sequencing** | | **Measures** | **C. elegans stage** | **Accession number and bigWig file** |
|---|---|---|---|---|
| ATAC-seq | | Chromatin accessibility | L1 | GSE114439_atac_wt_l1.bw |
| DNase-seq | | DNase I hypersensitivity | L1 | GSM3142662_dnase_wt_l1_rep1_100U_ml.bw |
| ChIP-seq | H3K4me1 | Histone modification (activation) | L1 | GSE114440_H3K4me1_wt_l1.bw |
| | H3K4me3 | Histone modification (activation) | L1 | GSE114440_H3K4me3_wt_l1.bw |
| | H3K27me3 | Histone modification (repression) | L1 | GSE114440_H3K27me3_wt_l1.bw |
| Short-capped RNA-seq | | Capped RNA 20-100 nt | L1 | GSE114490_scap_wt_l1_fwd.bw-GSE114490_scap_wt_l1_rev.bw |
| Long-capped RNA-seq | | Capped RNA > 200 nt | L1 | GSE114483_lcap_wt_l1_linear_fwd.bw-GSE114483_lcap_wt_l1_linear_rev.bw |
